# Developmental Manganese Exposure Causes Lasting Attention Deficits Accompanied by Dysregulation of mTOR Signaling and Catecholaminergic Gene Expression in Brain Prefrontal Cortex

**DOI:** 10.1101/2023.07.16.549215

**Authors:** N. A. Santiago, B. He, S. L. Howard, S. Beaudin, B. J. Strupp, D. R. Smith

## Abstract

Elevated manganese (Mn) exposure is associated with attentional deficits in children, and is an environmental risk factor for attention deficit hyperactivity disorder (ADHD). We have shown that developmental Mn exposure causes lasting attention and sensorimotor deficits in a rat model of early childhood Mn exposure, and that these deficits are associated with a hypofunctioning catecholaminergic system in the prefrontal cortex (PFC), though the mechanistic basis for these deficits is not well understood. To address this, male Long-Evans rats were exposed orally to Mn (50 mg/kg/d) over PND 1-21 and attentional function was assessed in adulthood using the 5-Choice Serial Reaction Time Task. Targeted catecholaminergic system and epigenetic gene expression, followed by unbiased differential DNA methylation and gene regulation expression transcriptomics in the PFC, were performed in young adult littermates. Results show that developmental Mn exposure causes lasting focused attention deficits that are associated with reduced gene expression of tyrosine hydroxylase, dopamine transporter, and DNA methyltransferase 3a. Further, developmental Mn exposure causes broader lasting methylation and gene expression dysregulation associated with epigenetic regulation, inflammation, cell development, and hypofunctioning catecholaminergic neuronal systems. Pathway enrichment analyses uncovered mTOR and Wnt signaling pathway genes as significant transcriptomic regulators of the Mn altered transcriptome, and Western blot of total, C1 and C2 phospho-mTOR confirmed mTOR pathway dysregulation. Our findings deepen our understanding of the mechanistic basis of how developmental Mn exposure leads to lasting catecholaminergic dysfunction and attention deficits, which may aid future therapeutic interventions of environmental exposure associated disorders.

**Significance Statement:** Attention deficit hyperactivity disorder (ADHD) is associated with environmental risk factors, including exposure to neurotoxic agents. Here we used a rodent model of developmental manganese (Mn) exposure producing lasting attention deficits to show broad epigenetic and gene expression changes in the prefrontal cortex, and to identify disrupted mTOR and Wnt signaling pathways as a novel mechanism for how developmental Mn exposure may induce lasting attention and catecholaminergic system impairments. Importantly, our findings establish early development as a critical period of susceptibility to lasting deficits in attentional function caused by elevated environmental toxicant exposure. Given that environmental health threats disproportionately impact communities of color and low socioeconomic status, our findings can aid future studies to assess therapeutic interventions for vulnerable populations.

## 1. Introduction

Attention and impulse control disorders, such as Attention Deficit Hyperactivity Disorder (ADHD), are the most prevalent neurodevelopmental disorders in children, affecting ∼6-11% of all U.S. children aged 6-17 years (1). Elevated environmental exposure to manganese (Mn) from contaminated well-water, ferroalloy industry emissions, and fungicide/pesticide use is associated with ADHD-like symptoms in children (2–18). The primary brain regions known to regulate attentional function are the prefrontal cortex (PFC) and striatum (19–21). The fronto-cortico-striatal brain circuit is believed to regulate attention through the catecholaminergic system, which is composed of neurotransmitters and receptors, such as dopamine and norepinephrine (19–21). Animal model studies have been important in establishing a link between developmental Mn exposure, lasting impairments in attention, impulse control, and sensorimotor function, and lasting hypofunctioning of the PFC and dorsal striatum catecholaminergic system (22–26). These Mn-associated effects include altered tyrosine hydroxylase (TH), dopamine transporter (DAT), dopamine D1 and D2 receptors (D1R, D2R), and norepinephrine transporter (NET) protein levels, and decreased K^+^-evoked release of dopamine and norepinephrine (22, 27-29) in fronto-cortico-striatal brain regions. However, the molecular mechanisms leading to these developmental Mn-induced catecholaminergic system changes are not well-understood.

Mn-induced alterations of the epigenome and transcriptome may occur due to the role of Mn as a pro-oxidant that can increase oxidative stress levels (30–32). Increased oxidative stress can increase DNA oxidation (e.g., as 8-OHdG and 5-hmc) to induce DNA hypomethylation, or produce elevated prooxidants that can alter DNA methyltransferase (DNMT) protein expression and form DNMT-containing complexes that facilitate DNA hypermethylation (30–32). It is known that DNA methylation of cytosine residues by DNMTs and histone deacetylation by hypoacetylhistone deacetylases (HDACs) regulate expression of key catecholaminergic genes TH, D1R, D2R, DAT, and NET (32–49). However, there is limited data on whether developmental Mn exposure may alter the epigenetic profile of catecholaminergic system genes (49–53). One example is that male mice exposed to perinatal Mn had hypermethylation of 24 gene promoter regions and transcript downregulation lasting into adulthood (51).

Recent evidence suggests that dysregulation of the neurodevelopmental signaling pathways mammalian/mechanistic target of rapamycin (mTOR) and Wingless-Int (Wnt) may lead to a hypofunctioning catecholaminergic system and an ADHD-like behavioral phenotype similar to that of our developmental Mn exposure rat model. For example, Wnt inhibition has been shown to decrease dopaminergic cell differentiation, myelination, dendrite morphogenesis, and activate mTORC1 signaling (54–61). Long lasting elevation in mTORC1 signaling is known to block canonical D1R signaling that is dependent on DARPP-32 (dopamine- and cAMP-regulated neuronal phosphoprotein) function (54–58). Further, mTORC1 activation can alter DNA methylation gene expression regulation through regulating DNMT1 and DNMT3A gene and protein expression (59–61). However, it remains unclear whether developmental Mn exposure may alter DNA methylation of mTOR and Wnt pathway genes, and whether these changes are associated with altered catecholaminergic system gene and protein expression levels.

Here, we sought to determine whether lasting attention deficits caused by developmental Mn exposure are accompanied by DNA methylation and gene expression changes in PFC genes in categories of epigenetic regulation, inflammation, cell development, and catecholaminergic system functions, including the Wnt and mTOR signaling pathways. Given that our observed Mn exposure-induced deficits persist into adulthood long after developmental Mn exposure has ended, we postulate that developmental Mn exposure produces lasting epigenome and gene expression changes of key PFC catecholaminergic and broader neuronal system functions. Male Long-Evans rats were exposed orally to Mn over PND 1-21 and attentional function was tested in adulthood using the 5-Choice Serial Reaction Time Task. Targeted gene expression analyses of the catecholaminergic system and epigenetic regulation genes, followed by unbiased differential DNA methylation, transcriptomics, and Western blot analyses were performed in young adult littermates. Our findings show that developmental Mn exposure causes lasting attention deficits that are accompanied by differential methylation and gene expression of inflammation, epigenetic regulation, cell development, and hypofunctioning neuronal systems. Furthermore, mTOR and Wnt pathways are identified as causal and upstream regulators of our Mn-altered transcriptome, and altered mTOR pathway function was evidenced by increased mTORC1 protein levels. These findings allow us to propose a set of novel comprehensive mechanisms for how developmental Mn exposure leads to lasting catecholaminergic dysfunction and attention deficits, which may aid future therapeutic interventions of environmental exposure-associated neurobehavioral disorders.

## 2. Methods

### 2.1. Subjects

Subjects were 128 neonate male Long-Evans rats born in-house from nulliparous timed-pregnant females (from Charles River gestational day 18). Twelve to 24 hours after parturition (designated PND 1, birth = PND 0), litters were sexed, weighed, and culled to eight pups per litter such that each litter was composed of five to six males and the remainder females. Only one male per litter was assigned to a particular Mn treatment condition. Rats (dams and weaned pups) were fed Harlan Teklad rodent chow #2920 (reported by the manufacturer to contain 80 mg Mn/kg) and housed in polycarbonate cages at a constant temperature of 21 ± 2°C. At PND 22, all pups were weaned and the rats designated for behavioral testing were pair-housed with an animal of the same treatment group and maintained on a reversed 10:14 hr light/dark cycle. All aspects of behavioral testing and feeding were carried out during the active (dark) phase of the rats’ diurnal cycle. Littermates that were designated for molecular analysis were grouped by treatment and fed *ad libitum* until PND 66 when sacrifice and brain tissue collection was performed for molecular analysis.

Males were exclusively used because attentional dysfunction is two to three times more prevalent in boys than girls (62–63), and because our prior studies have established that early postnatal Mn exposure causes lasting impairments in learning, attention, and fine motor function in male rats (22–29). All animal procedures were approved by the institutional IACUC (protocols Smithd1803) and adhered to National Institutes of Health guidelines set forth in the Guide for the Care and Use of Laboratory Animals. Criteria for exclusion of rats from the study were based on overt signs of poor rat health, including loss of body weight, absence of grooming, impaired function, and death; no rats were excluded from the study based on these criteria. This study was not pre-registered.

### 2.2. Manganese dosing regimen

Neonatal rats were orally exposed to a Mn dose of 50 mg Mn/kg/day starting on PND 1 through weaning on PND 21 (early postnatal Mn exposure) (Supplemental Methods 2.2). This Mn exposure regimen is relevant to children exposed to elevated Mn via drinking water and/or diet; pre-weaning exposure to 50 mg Mn/kg/day produces a relative increase in Mn intake about the level of increase reported in infants and young children exposed to Mn-contaminated water or soy-based infant formulas (22–29).

### 2.3. Focused attention

The 5-Choice Serial Reaction Time Task (5-CSRTT) was used to assess attentional function as previously described (24–26). Briefly, rats began testing at about PND 45 (60-62 rats/treatment), with food magazine and nose-poke training for 1 week, followed by two 5-choice visual discrimination tasks using fixed cue durations of 15 and 1 second, respectively. Once the animals attained 80% correct on two of three successive sessions in the visual discrimination tasks, they moved on to the focused attention tasks. Two successive focused attention tasks were used to assess the ability of the rats to detect and respond to a brief visual cue presented unpredictably in time and location (i.e., one of the five response ports). The first focused attention task was administered over PND 73–86 and used pre-cue delays of 0, 1, 2, and 3 seconds. The second and more challenging focused attention task was conducted over PND 88–102, and used pre-cue delays of 0, 3, 4, and 5 seconds (Supplemental Methods). All rats were weighed and tested 6 days/week throughout training and testing. Behavioral assessment occurred during the active (dark) period of the diurnal cycle at the same time each day and in the same operant chamber for each individual rat. All behavioral testing was conducted by individuals blind to the treatment condition of the subjects. All rats were maintained on a food restriction schedule with water available *ad libitum* throughout behavioral assessment, as described previously (23–26).

### 2.4. Targeted catecholamine and epigenetic system gene expression analysis

Gene expression of key catecholaminergic (*Th*, *Dat*, *Drd2*) and epigenetic modulator genes (*Dnmt1*, *Dnmt3a*, *Dnmt3b*, *Hdac3*, and *Hdac4*) was determined through RT-qPCR on PFC tissue from PND 66 control and developmental Mn-exposed littermates of the behaviorally tested rats (6-7 rats/treatment). This age was selected because it coincides with the age at the start of behavioral testing in the focused attention tasks (22–29). Briefly, partially thawed fresh-frozen PFC tissue punches (1.39 mm thickness, 1.5 mm diameter, 25-30 mg) were taken from the anterior cingulate cortex/prefrontal cortex area (Paxinos and Watson Rat Brain Atlas: 2.16-0.77 mm anterior to bregma) over dry-ice and underwent Dounce Homogenization and total RNA extraction, DNAse treatment, and Reverse Transcription. RT-qPCR cycle quantity thresholds were measured using the ThermoFisher Scientific TaqMan Advanced Master Mix (Applied Biosystems: Cat. #4444556) and TaqMan primers on the Bio-Rad CFX95 instrument (Supplemental Table 1) following the manufacturer’s protocol. *Gapdh*, *ActB*, and *Ubc* expression were assessed as reference genes and the tool Normfinder confirmed the most stable reference gene combination for gene expression analysis (64–65). RT-qPCR was performed with two RT reaction replicates per animal, and three replicates per RT for each animal (66) (Supplemental Methods).

### 2.5. Differential DNA methylation and functional pathway analysis

We conducted reduced representation bisulfite sequencing (RRBS) to determine whether developmental Mn exposure led to lasting alterations of genome methylation status that may associated with the n-induced hypofunctioning catecholaminergic system identified in our prior studies (22–29). DNA was extracted from the same PND 66 fresh-frozen PFC samples detailed in Section 2.4, bisulfite converted, and sequenced at the University of California Davis Genome Center (n=3 rats/treatment). Galaxy pipeline analysis was conducted to obtain differentially methylated regions (DMRs) between control and Mn groups. Significant DMRs (unadjusted and adjusted p<0.05) were annotated by chromosome to the rat (rn6) genome, and then visualized and manually confirmed using the Integrative Genomics Viewer (IGV) sliding window (67). DMR fold-change values were calculated for each differentially methylated gene as the difference of the mean methylation value of the control versus Mn PFC samples. All significant (p<0.05) DMR genes were then ranked and analyzed using Fast Preranked Gene Set Enrichment Analysis (FGSEA) to determine the associated functions of the differentially methylated gene products through Gene Ontology biological processes (68). Gene Ontology biological processes groups were further reduced, filtered, and visualized using REVIGO ontology’s algorithm for semantic similarity of parent and child terms (69). Additionally, Kyoto Encyclopedia of Genes and Genomes (KEGG) analysis were conducted on all significant DMR associated genes (p < 0.05 unadjusted) using EnrichR (70) (Supplemental Methods).

### 2.6 Differentially expressed genes and functional pathway analysis

The ability of developmental Mn exposure to alter global gene expression was determined by 3’ Tag RNA-sequencing. RNA aliquots from the same PND 66 PFC RNA extraction in Method 2.4 (n=5-6 rats/treatment) were analyzed for Bioanalyzer quality assessment, library preparation, and 3’ Tag RNA-sequencing by the University of California Davis Genome Center. FastQ files were processed using established Geneious Prime RNA-sequencing tools (Geneious Prime 2022.0). The tool DESeq2 was used to calculate the fold-change and p-value significance of the developmental Mn-exposed PFC differentially expressed genes (DEG), relative to control rats (71). Next, all DEGs were ranked and analyzed using Fast Preranked Gene Set Enrichment Analysis (FGSEA) to determine the DEG’s Gene Ontology-associated biological processes, molecular functions, and cellular components subtypes. Gene Ontology biological processes groups were further reduced, filtered, and visualized using REVIGO’s algorithm for semantic similarity of parent and child terms (69). The reduced Gene Ontology terms were then categorized into the four functional categories of inflammation, epigenetics, cell development, and neuronal function, based on our *a priori* hypotheses and known mechanisms of Mn neurotoxicity (22, 27-30, 49-52, 72). KEGG analysis was conducted on all significant DMR associated genes (p < 0.05 unadjusted) using EnrichR (70)(Supplemental Methods).

### 2.7. Differential methylation and differential expression integration

Integrated analysis of DMR and DEG data was conducted to determine the Gene Ontology biological processes associated with both differential methylation and differential gene expression of our Mn neurotoxicity attention deficit phenotype. We used the Genomics Tools Venn-Diagram web program and REVIGO to compare and visualize the upregulated and downregulated Gene Ontology biological processes group lists shared by both DMR associated genes and DEGs, and the four *a priori* hypothesized categories of biological dysfunction (inflammation, epigenetics, cell development, and neuronal function) described in Methods 2.5. Genomics Tools Venn-Diagram web program comparison was performed on the differentially methylated and expressed genes to further determine genes that were both differentially methylated and differentially expressed. Clustergrammer was then used to visually display DMR and DEG expression levels, and the gene region locations of the DMRs (e.g., promoter, exon, intron, etc.) were determined using the UCSC Genome Integrator and IGV slide sorting noted in Methods 2.5.

Ingenuity Pathway Analysis (IPA) was performed at the UCLA Technology Center for Genomics & Bioinformatics using all p-unadjusted DEGs from DESeq2 to identify significant causal and upstream regulator genes. Genes that were identified as upstream and causally-related to the DEGs were then integrated with the differentially methylated and expressed genes using the Genomics Tools Venn-Diagram web program comparison to identify genes that may be mechanistically associated with the Mn-neurotoxicity DEG phenotype. Finally, in order to further strengthen our assessment of the significance of the identified causal and upstream regulators, we compared the DMR and DEG lists to the most significant activation value regulator Search Tool for the Retrieval of Interacting Genes/Proteins (STRING) gene interaction pathway lists (73)(Supplemental Methods).

### 2.8. Total and phosphorylated mTOR protein levels

Western blots were conducted for total mTOR, mTORC1, mTORC2, and β-Catenin protein levels to determine whether developmental Mn exposure led to lasting dysfunction of the mTOR and Wnt signaling pathway. Briefly, protein was extracted from PFC brain punches of PND 66 control and Mn rats, homogenized, and protein lysates were prepared for Western blot analysis (Supplemental Table 2 for antibodies). Western blot band densitometry was quantified using Bio-Rad Image Lab volume quantification software (5-6 rats/treatment) (Supplemental Methods).

### 2.9. Mn Blood levels

Blood Mn concentrations were measured in PND 24 littermates and the same PND 66 rats used in the reported molecular outcomes using methods previously described (n=7 rats/treatment/age) (22–29). Mn levels were determined using a Thermo Element XR inductively coupled plasma–mass spectrometer. The analytical detection limit for Mn in blood was 0.04 ng/mL (Supplemental Methods).

### 2.10. Statistical analysis

The behavioral data were modeled by way of structured covariance mixed models. Fixed effects included in the model were Mn exposure (two levels corresponding to the control and Mn-treated groups), pre-cue delay, cue duration, and session block depending on the outcome analyzed. For analysis, the 12 days of testing for each focused attention task were designated as four 3-day test session blocks. Statistical tests used a Kenward-Roger correction. Plots of residuals by experimental condition were used to examine the assumption of homogeneity. Additional random effects with high variance in the residuals across levels of the factor (e.g., session block) were added to achieve homogeneity. The distribution of each random effect was inspected for approximate normality and presence of outliers. The significance level was set at p ≤ 0.05. Significant main effects or interaction effects were followed by single-degree of freedom contrasts to clarify the nature of the interactions, using the Student’s t-test for pairwise comparisons of least squared means. All behavioral data analyses were conducted using SAS (version 9.4) for Windows. RT-qPCR gene expression data statistical analyses were performed on ΔCt values by non-parametric Wilcoxon rank-sum/Kruskall-Wallis test analysis and p ≤ 0.05 deemed significance (74). Western blot protein expression data were analyzed by non-parametric Wilcoxon rank-sum/Kruskall-Wallis test to compare two paired groups, with significance set at p ≤ 0.05. Mn blood levels were analyzed by ANOVA and Tukey-Post hoc analysis to establish p ≤ 0.05 statistical significance. These latter analyses were performed using JMP Pro (version 16.1.0; SAS Institute, Inc.).

## 3. Results

### 3.1. Developmental Mn exposure impairs focused attention

We performed two successive focused attention tasks using the 5-CSRTT to test whether developmental Mn exposure causes lasting deficits in attentional function in these animals, consistent with our prior studies (25–26). Focused attention can be defined as the ability to maintain attentional focus to detect an unpredictable visual cue, in the face of variable delays between trial onset and presentation of the visual cue (25). The first focused attention task (FA1) used variable pre-cue delays of 0, 1, 2, or 3 and a fixed visual cue duration of 0.700 sec (See Methods 2.3.). The analysis of percent accurate responses revealed a significant interaction between Mn exposure and testing session block (F(3, 269.5)=4.33, p=0.005). Although the Mn group performed significantly more poorly than controls across all four test session blocks (p’s = 0.035, 0.002, <0.001, and < 0.001 for session blocks 1-4, respectively) (Figure 1A), the difference in performance between groups grew larger across testing blocks because the controls improved more across testing than the Mn group, based on within treatment group contrasts across session blocks (Figure 1A). In the second focused attention task (FA2), which is more challenging with variable longer pre-cue delays of 0, 3, 4, or 5 sec and visual cue durations of 0.4 or 0.7 sec, we found that the Mn-exposed animals continued to exhibit lower attentional accuracy compared to controls, based on a significant main effect of Mn exposure (F(1, 228)=9.7, p=0.002) (Figure 1B). However, there was no higher order interaction between Mn and block (F(3, 579)=1.73, p=0.1597), or between Mn and pre-cue delay (F(3, 508)= 0.50, p=0.685). Altogether these results establish that developmental Mn exposure causes lasting deficits in attention, consistent with our previous published studies (22, 25-26).

**Figure 1:**
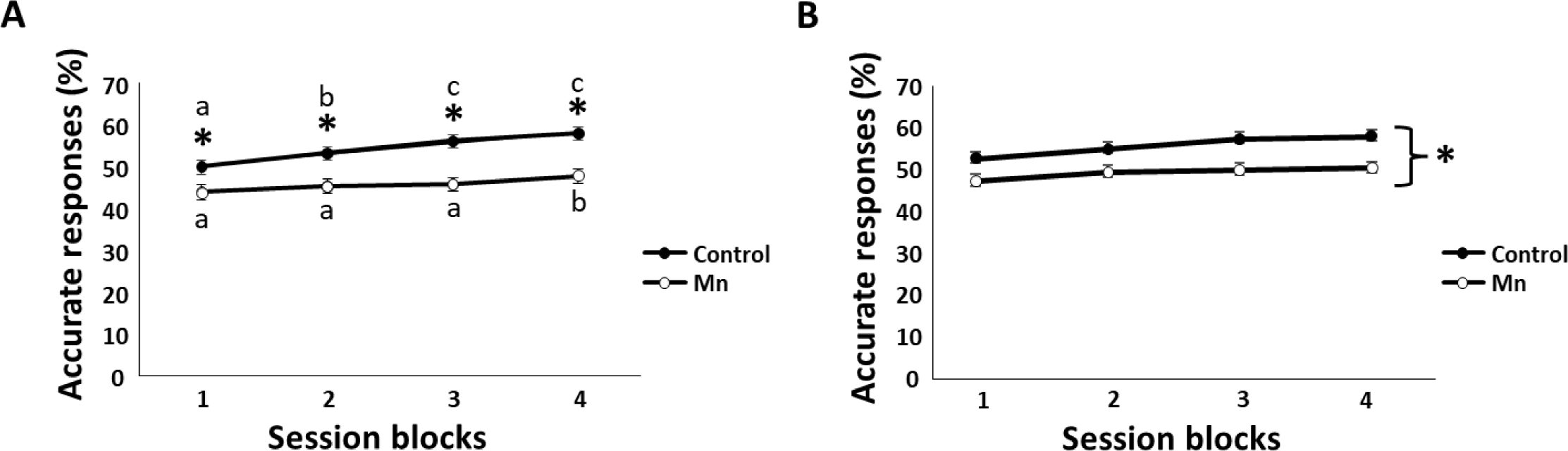
Developmental Mn exposure impairs focused attention. Percent accurate responses of the control and Mn groups in the first (A) and second (B) focused attention tasks as a function of testing session block (3 days of testing/block). * p < 0.05 versus controls in ‘A’ and ‘B’. Letter superscripts in ‘A’ indicate within treatment group differences (p < 0.05) between session blocks; within treatment group, session blocks with different superscript letters are significantly different (p < 0.05). Data are lsmeans ± SEM (n=60-62/treatment).

### 3.2. Developmental Mn exposure alters expression of key catecholaminergic system and epigenome associated genes

Protein levels of the key catecholaminergic system genes *Th*, *Dat*, and *Drd2,* which are known to play a role in attentional function, are also known to be altered by developmental Mn exposure (22, 27-29). Thus, we sought to assess whether developmental Mn exposure leads to lasting changes in the expression of these genes, as well as their known epigenetic gene expression regulators *Dnmt1*, *Dnmt3a*, *Dnmt3b*, *Hdac3*, and *Hdac4* in the prefrontal cortex (PFC) of young adult animals (32–49). Results show that developmental Mn exposure causes lasting ∼50% and ∼83% reductions in *Th* and *Dat* gene expression, respectively, relative to controls (p’s=0.05 and 0.03), while *Drd2* gene expression was not measurably affected (p=0.25) (Figure 2). Developmental Mn exposure also decreases *Dnmt3a* gene expression by 31% (p=0.05), but does not significantly change *Dnmt1*, *Dnmt3b*, *Hdac3*, or *Hdac4* gene expression levels (p’s = 0.10, 0.20, 0.32, and 0.31, respectively) (Figure 2). These findings demonstrate that developmental Mn exposure causes lasting alteration in gene expression of key catecholaminergic system genes, consistent with our prior studies showing changes in catecholaminergic system protein levels, and they suggest and a potential role for an epigenetic mechanism underlying our Mn exposure-associated attentional deficits (22, 27–29).

**Figure 2:**
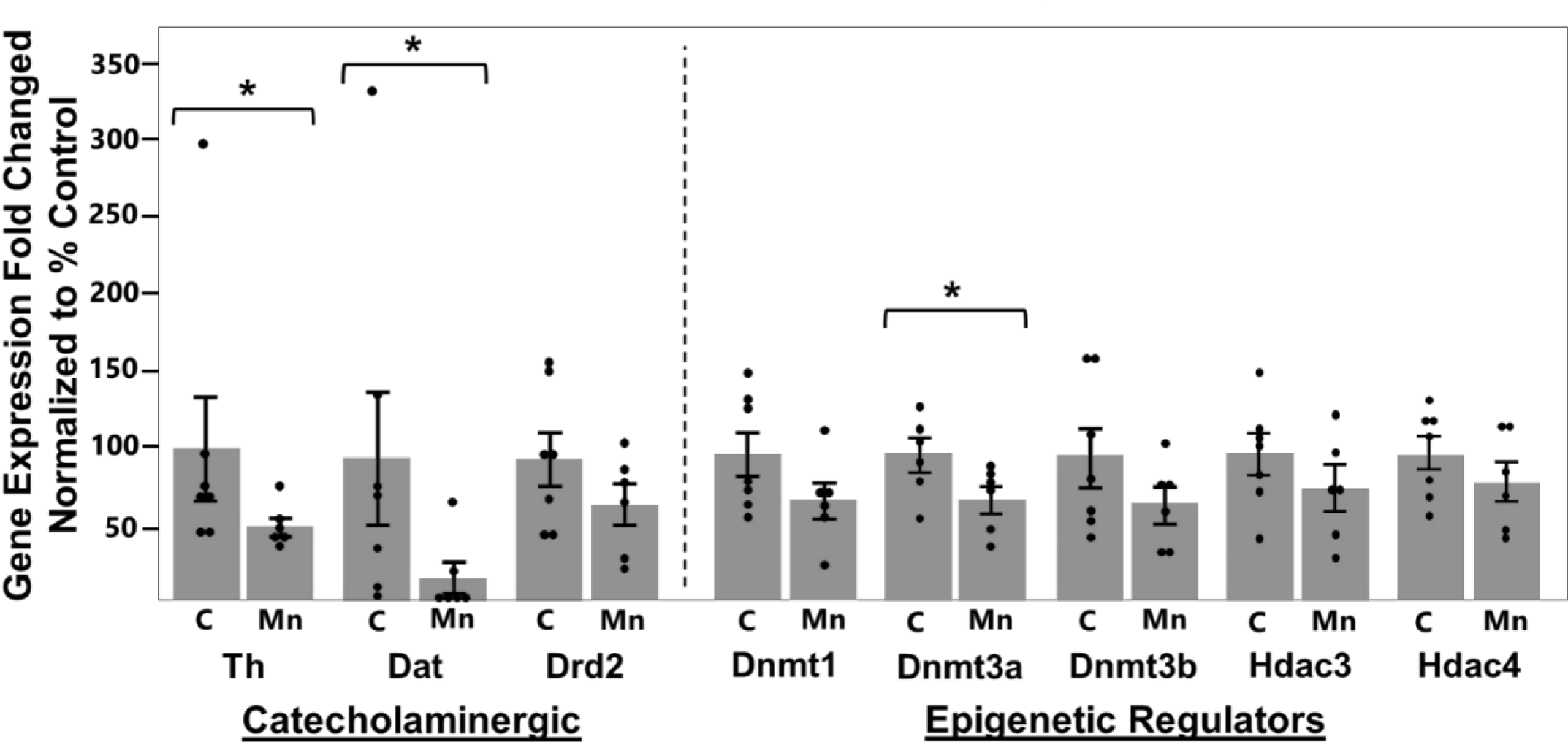
Developmental Mn exposure alters expression of key catecholaminergic and epigenetic genes. PND 66 prefrontal cortex RT-qPCR gene expression results conducted on catecholaminergic and epigenetic modulator genes. Gene expression fold-change values are presented as normalized to % of control (all ΔΔCt values divided by the mean of control and multiplied by 100). Pair-wise statistical comparisons were performed on ΔCt values. * indicates significant p < 0.05 versus control, based on non-parametric Wilcoxon analysis (n=6-7 rats/treatment). Error bars represent SEM.

### 3.3. Developmental Mn exposure leads to global hypermethylation of genes associated with inflammatory, epigenetic, cell development, and neuronal system changes in the PFC

We conducted reduced representation bisulfite sequencing (RRBS) to determine whether developmental Mn exposure leads to lasting alterations of PFC genome methylation status that may contribute to the Mn-induced hypofunctioning catecholaminergic system. Overall, Mn exposure induces differentially methylated regions (DMRs) associated with 4250 genes (p-unadjusted < 0.05), of which ∼83% (i.e., 3541/4250) are hypermethylated and 17% (709/4250) hypomethylated. After adjusting for multiple comparisons, 1027 genes remain hypermethylated and 192 genes remain hypomethylated (p-adjusted < 0.05) (Figure 1A; Supplemental Sheet 1). In regard to our hypothesis that developmental Mn exposure causes lasting disruption of epigenetic and catecholaminergic system gene methylation state, specifically, we found that the epigenetic modulators DNA methyltransferase (1, 3a, and 3b) and histone deacetylase (4 and 7) are hypermethylated, while histone deacetylase 3 is hypomethylated (Supplemental Sheet 1). Moreover, several catecholaminergic system and Mn transport-related genes are differentially methylated, including hypermethylation of dopamine receptor D4 (*Drd4*), tyrosine 3-monooxygenase/tryptophan 5-monooxygenase activation protein theta (*Ywhaz*), catechol-O-methyltransferase (*Comt*), solute carrier family 30 member 10 (*Slc30a10*), and protein phosphatase 1 regulatory inhibitor subunit 1B (*Ppp1r1b*/DARPP-32), and hypomethylation of 3-monooxygenase/tryptophan 5-monooxygenase activation protein gamma (*Ywhag*) and GABA transporter 1 (*Slc6a1*) (p-adjusted p<0.05) (Supplemental Sheet 1).

Gene Ontology biological processes analyses were conducted on all significant 4250 DMR genes (p < 0.05 unadjusted) to elucidate the overall biological processes associated with our Mn-induced differentially methylated genes. The results identify 814 hypermethylated and 239 hypomethylated Gene Ontology terms (p < 0.05), many of which align with our *a priori* hypothesized mechanistic categories of Mn neurotoxicity (i.e., alteration of inflammation, epigenetic regulation, cell development, and neuronal function-related processes) (Figure 1B; Supplemental Sheet 2). Of particular relevance to pathways known to regulate catecholaminergic system function is the identification of the hypermethylated *(GO:0060070) canonical Wnt signaling pathway* and the hypomethylated *(GO:0032008) positive regulation of TOR signaling* (Figure 1B). Subsequent KEGG pathway analysis revealed that the differentially methylated genes are associated with neurodegenerative disease and that the *mTOR signaling pathway* is a top hypermethylated pathway caused by developmental Mn exposure (Supplemental Figure 1).

### 3.4. Developmental Mn exposure induces lasting gene expression changes associated with altered inflammatory, epigenetic, cell development, and neuronal system changes in the PFC

To assess whether broader gene expression changes could be associated with the differential methylation changes noted above and inform the mechanistic basis for the lasting deficits in attentional function, we conducted RNA-seq analysis on PFC brain tissue. Findings show developmental Mn exposure leads to lasting differential expression of 501 (p-unadjusted < 0.05) or 27 (p-adjusted < 0.05) genes (Figure 4A; Supplemental Sheet 3). Of the 27 p-adjusted differentially expressed genes, 13 are upregulated and 14 are downregulated (Figure 4A). All 27 genes can be readily categorized into the four *a prior* hypothesized biological function categories identified in the differential methylation analyses above using the Rat Genome Database, as follows: inflammation (eight genes), epigenetics (five genes), cell development (four genes) and neuronal function (10 genes) (Figure 4A). Notably, several of the p-adjusted differentially expressed genes are associated with catecholaminergic function and Wnt and mTOR signaling pathways. Specifically, the encoded proteins of *Crebbp*, *Prkacb*, *Gabarapl1*, and *Ywhaz* are known regulators and intermediates of the Wnt and mTOR pathway regulating neuronal cell proliferation and differentiation, and specifically, their gene and protein expression may alter dopamine synthesis, release, and transport pathways (Supplemental Sheet 4).

Gene Ontology biological processes enrichment analysis was conducted to gain further insight into broader gene expression pathway changes caused by developmental Mn exposure. Results identify 192 down- and 211 up-regulated (p-unadjusted < 0.05) Gene Ontology biological processes terms (Supplemental Sheet 5). These Gene Ontology terms were further refined, filtered, and visualized using REVIGO, leading to identification of 69 down- and 63 up-regulated terms that were then categorized into the four *a priori* hypothesized mechanistic categories noted above (i.e., inflammation, epigenetic regulation, cell development, and neuronal function), and their respective leading edge contributing genes as defined by Gene Ontology enrichment analysis and fold-changes were determined. Consistent with our hypothesis, one notable neuronal function term of relevance that has genes with reduced expression is *(GO:0033605) positive regulation of catecholamine secretion*, while cell development terms with genes with increased expression are *(GO:0031929) TOR signaling* and *(GO:0030178) negative regulation of Wnt signaling* (Figure 4B, C; Supplemental Figure 4). Additional Gene Ontology molecular function and cellular component and KEGG pathway analyses illustrates that developmental Mn exposure is associated with lasting proinflammatory and hypofunctioning neuronal systems (Supplemental Figure 2, 3). Overall, the Gene Ontology differential gene expression analysis further supports a significant pattern of mTOR, Wnt, and hypofunctioning neuronal pathways caused by developmental Mn exposure.

### 3.5. Integration of Mn induced differential methylation and gene expression further supports mTOR and Wnt pathways dysregulation is associated with hypofunctioning neuronal systems

We conducted an integrated analysis of the Gene Ontology terms shared by both differentially methylated and differentially expressed genes to evaluate whether there were altered neuronal functions that may inform our Mn attentional deficit phenotype. We found that 27% (41/148) of our downregulated gene expression-associated Gene Ontology terms are hypermethylated, while about 7% (14/204) of our upregulated Gene Ontology terms are hypomethylated (Figure 5A). In a Venn diagram comparison of all significant Gene Ontology terms we see that the *(GO:0032006)* regulation of *mTOR signaling* and *(GO:0035567) non-canonical Wnt signaling* pathways are present in both hypermethylated and hypomethylated Gene Ontology term lists and the Gene Ontology term *(GO:0010467) gene expression* is present in the hypermethylated, gene expression upregulated, and gene expression downregulated Gene Ontology term lists (Figure 5B). Overall, DMR and DEG integrated Gene Ontology analysis shows that developmental Mn exposure is associated with mTOR and Wnt signaling dysregulation that is also associated with genes and pathways that regulate gene expression, as determined by the shared Gene Ontology terms.

Subsequently, we assessed which genes of our DMR and DEG analysis were both differentially methylated and expressed and we found that developmental Mn exposure leads to the lasting differential methylation and expression of 155 genes in the PFC (p < 0.05-adjusted) (Figure 5C, D; Supplemental Sheet 6). Notably, of these 155 genes, 85% are differentially methylated within their promoter region, while the remaining 15% are differentially methylated across various exons or their terminator region (Supplemental Sheet 6). Gene Ontology biological function, molecular function, and cellular components analysis of the 155 differentially methylated and differentially expressed genes were easily categorized into the same inflammatory, epigenetic, cell developmental, and hypofunctioning neuronal pathways functional categories described above (Figures 3, 4; Supplemental Figure 5). Also of note is that subsequent KEGG pathway analysis using the 155 differentially methylated and expressed genes identified the *mTOR signaling pathway* as the most significant upregulated term by p-value (p = 0.003), along with several downregulated neuronal pathways, such as the *GABAergic synapse* (Supplemental Figure 6). Altogether, the functions of the 155 differentially methylated and expressed genes support our hypothesis that developmental Mn exposure leads to lasting mTOR signaling dysregulation and hypofunctioning neuronal systems.

### 3.6. IPA causal and upstream regulator integration with DMRs and DEGs corroborate Wnt and mTOR pathways are novel mechanistic targets of developmental Mn exposure

We performed Ingenuity Pathway Analysis (IPA) on the 501 (p-unadjusted < 0.05) genes that emerged from the DESeq2 analysis reported above, in order to determine whether causal and upstream regulator gene(s) may help explain the lasting gene expression phenotype caused by developmental Mn exposure. IPA causal analysis reveals 447 (p-unadjusted < 0.05) significant genes, while IPA upstream analysis reveals 179 (p-unadjusted < 0.05) genes that are associated with (i.e., regulate) our transcriptomic findings (Supplemental Sheet 7, 8). Of these, 129 genes are identified as both upstream and causal regulator genes, and eight of these genes (*Dnajc5*, *Qki*, *Lrrc4*, *Immt*, *Chchd6*, *Gabbr2*, *Prkacb*, *Uchl3*) are themselves differentially expressed (p-unadjusted) (Figure 6B; Supplemental Figure 7). The top 15 significant causal and upstream regulator genes, based on their IPA-predicted activation value, are shown in Figure 6A, along with a brief description of their function. Notably, these regulator genes are also readily categorized within the four *a priori* hypothesized mechanistic categories of Mn neurotoxicity noted above (inflammation, epigenetic regulation, cell development, and neuronal function), and they share several of the same Gene Ontology biological process and KEGG terms from the DMR and DEG analysis (Figures 3-5; Supplemental Figures 1-6).

**Figure 3:**
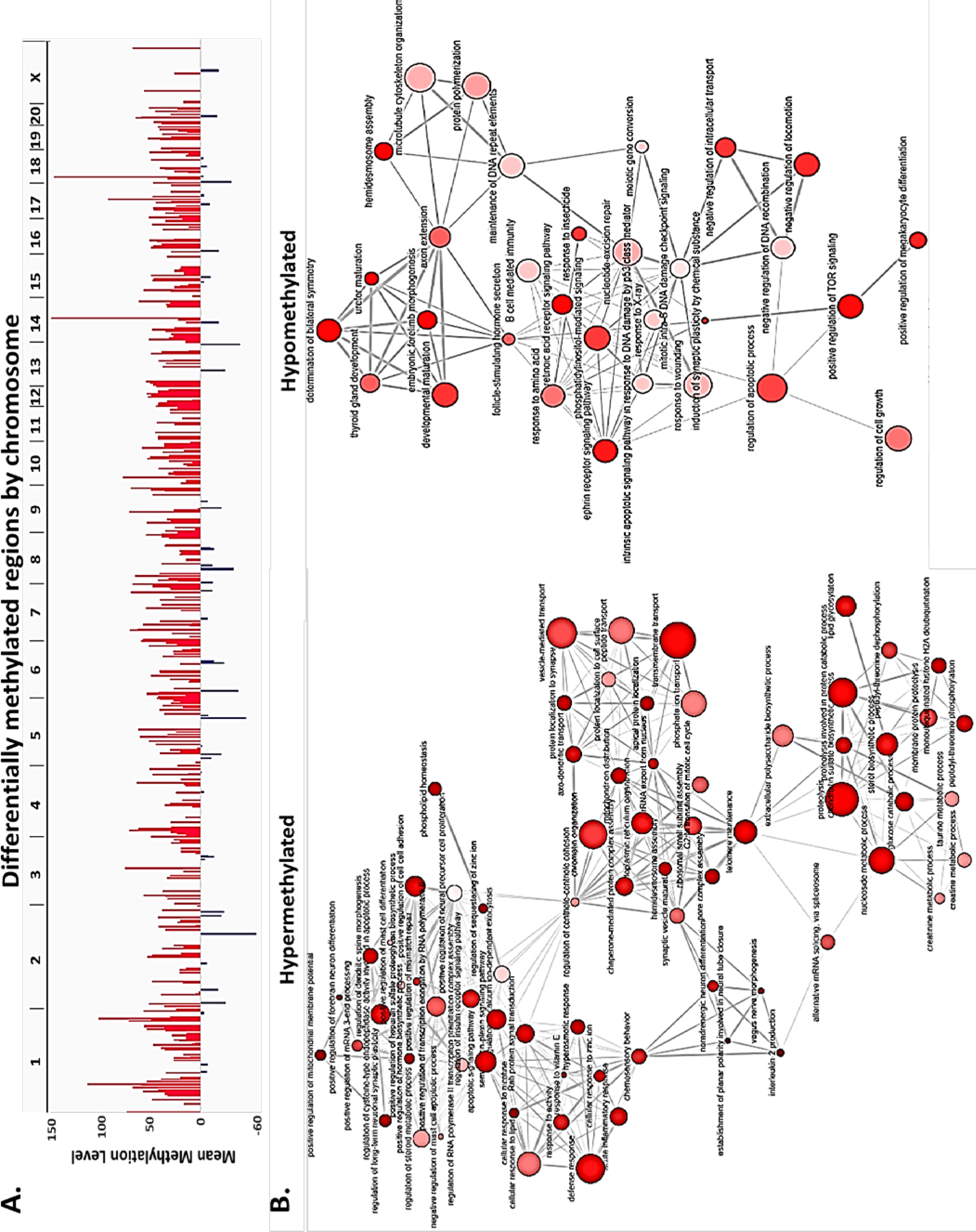
Developmental Mn exposure leads to differential methylation in inflammatory, epigenetic regulation, cell development, and neuronal systems. (A). RRBS of PND 66 PFC tissue, all significant (p-adj.<0.05) differentially methylated regions. Red=hyper-/Blue=hypo-methylated, mean methylation levels. (B). All p < 0.05 unadjusted Gene Ontology term similarity connections maps. All circles are p < 0.05 significant GO terms. Smaller circle = smaller p-val/Darker = more child terms included in parent term. Thicker connecting line between terms indicates more child terms within that share a functional connection. Figure is shown on separate page below.

**Figure 4:**
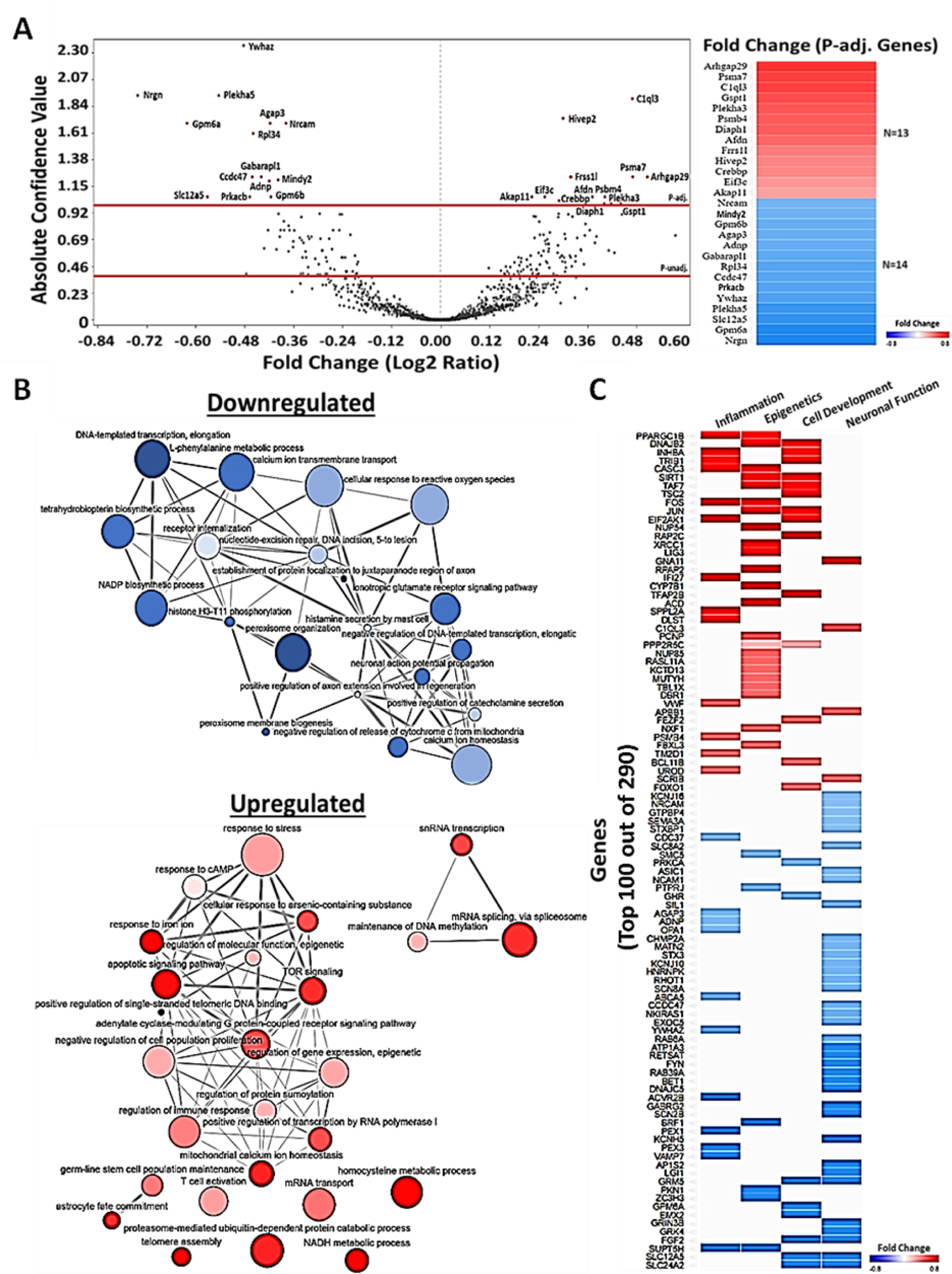
Developmental Mn exposure induces lasting gene expression changes associated with altered inflammatory, epigenetic regulation, cell development, and neuronal system biological function in the PFC. (A). Fold change expression levels of developmental Mn exposure induced differentially expressed genes (DEGs) are displayed as a volcano plot. Each red line is marked as the genes that are significant before and after multiple comparison adjustments. (B). The list of p-adjusted DEGs are displayed on the heatmap on the right by their respective fold changes. (B). Gene Ontology similarity connections maps are displayed by downregulated or upregulated genes that contributed to the Gene Ontology term results. All circles represent p < 0.05 GO terms. Smaller circle size=smaller p-value/Darker=more child terms that have been reduced to the parent term represented. Thicker connecting line between two Gene Ontology terms indicates more child terms within both terms. (C). A heatmap displays the leading edge DEGs and their log2 Fold Change levels of the categorized (inflammation, epigenetic, cell development, and neuronal function) Gene Ontology terms. Figure is shown on separate page below.

**Figure 5:**
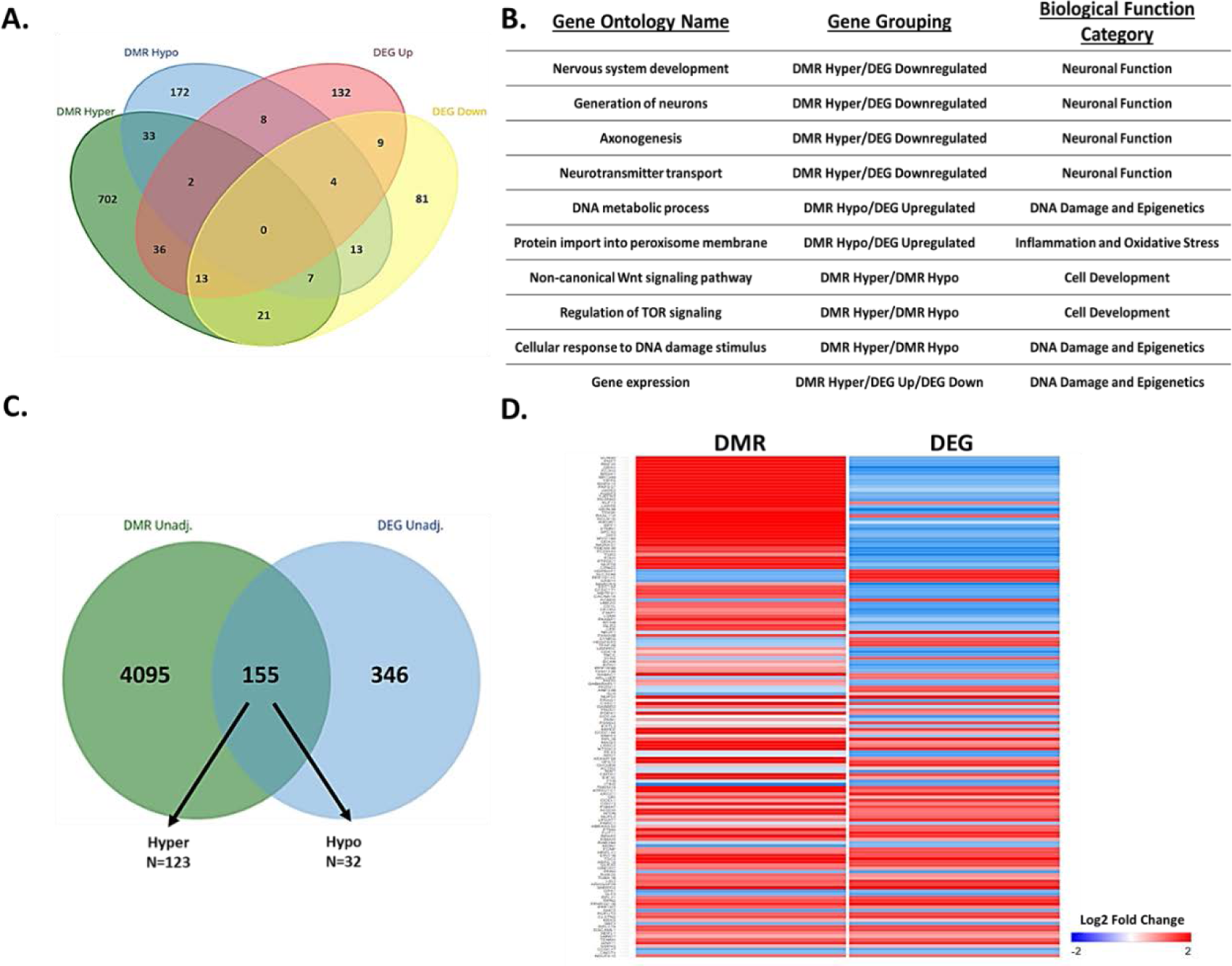
Mn exposure led to 155 PFC genes being differentially methylated and expressed in functions related to hypofunctioning neuronal systems. (A). Venn diagram of shared Gene Ontology groups between DMR hyper-and hypomethylated associated genes and DEG up-and downregulated genes. (B). Table of top relevant Gene Ontology groupings and their biological category significance. (C). DMR and DEG genes are compared to identify which genes are both differentially methylated and differentially expressed. (D). The 155 genes that are both DMR and DEG are displayed in a heatmap with their corresponding fold change values (See Supplemental Sheet 6 for clearer readability of genes in ‘D’).

**Figure 6:**
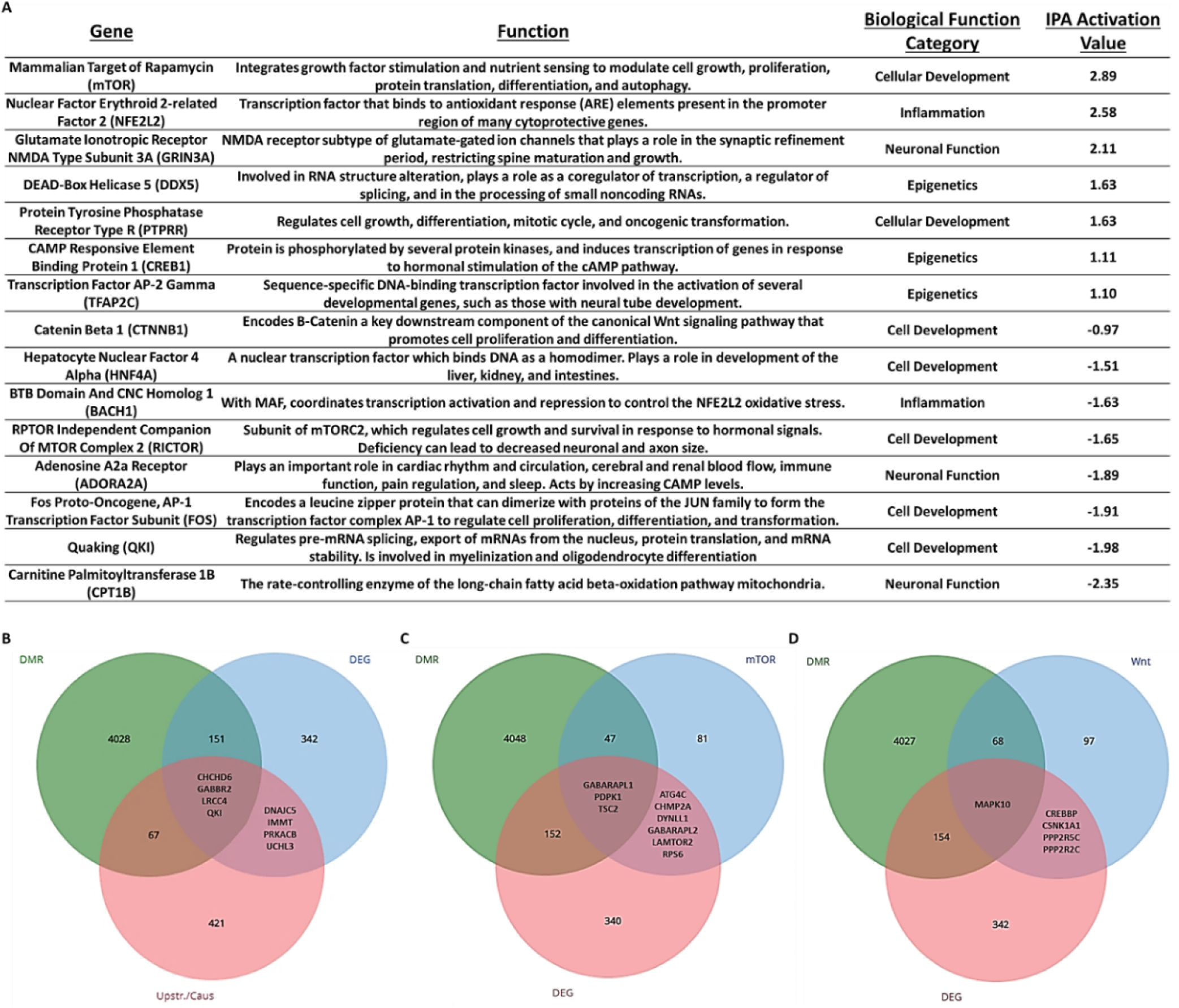
Integration of IPA causal and upstream regulators with significant DMRs and DEGs elucidates. (A). The top 15 (by activation value) significant upstream and causal gene regulators determined by Ingenuity Pathway Analysis (IPA) are displayed with a brief definition of biological relevance to our Mn neurotoxicity phenotype, the biological function category that matches the Gene Ontology Group categories, and the IPA activation value. IPA activation values above 2 and below −2 are predicted statistically to be activated in their corresponding upregulated or downregulated direction. (B). A Venn diagram of the resulting integration of IPA Causal and Upstream Regulators with DMR and DEGs. (C). Known genes regulated by mTOR compared to the significant Mn altered DMR and DEGs. (D). Known genes regulated by Wnt compared to the significant Mn altered DMR and DEGs. mTOR and Wnt pathway dysregulation as key Mn mechanistic targets.

It is highly noteworthy that the mTOR signaling pathway is consistently identified as being altered by, as well as mediating many of the lasting molecular changes caused by developmental Mn exposure, as it has the largest IPA activation value and a high frequency of interactions (8/15 other top master regulators) defined in the gene interaction pathways (Figure 6A). The Ctnnb1/Wnt signaling pathway is also identified as a top regulator, with regulation interactions with several of the other top 15 regulators identified in the IPA analyses (Figure 6A). Together, the mTOR and Wnt master regulator gene interaction networks can account for (i.e., regulate) 15 and 32 of the 501 differentially expressed genes, respectively (Supplemental Figure 8).

Finally, to further strengthen our assessment of the significance of Wnt and mTOR regulatory pathways altered by developmental Mn exposure, we compared our lists of differentially methylated and expressed genes to the Search Tool for the Retrieval of Interacting Genes/Proteins (STRING) gene interaction pathway lists for both mTOR and Wnt signaling pathways (Figure 6C, D). This analyses shows that, of the 4250 p-unadjusted differentially methylated genes, 50 genes were determined through STRING to be associated with mTOR signaling pathways and 69 genes were associated with Wnt signaling pathways (Figure 6C, D; Supplemental Sheet 9). In particular, five differentially expressed genes were associated with Wnt STRING genes and nine differentially expressed genes were associated with mTOR STRING genes (Figure 6C, D; Supplemental Table 9). These results further highlight that developmental Mn exposure is associated with important lasting gene expression changes mediating neuronal function, particularly the Wnt and mTOR signaling pathways.

### 3.7. Developmental Mn leads to decreased total mTOR and increased mTORC1 protein levels in rat PFC, which establishes impaired mTOR signaling as a novel mechanistic target of Mn neurotoxicity

We conducted Western blot analysis for total mTOR, phosphorylated-mTORC1, phosphorylated-mTORC2, and β-Catenin (proxy for Wnt signaling) protein levels in brain PFC to determine whether the lasting differential methylation and gene expression changes caused by developmental Mn exposure reported above translated to functional dysregulation of the mTOR and Wnt signaling pathways. We found that developmental Mn exposure leads to a 28% decrease in total mTOR (p = 0.008) and a 51% increase in mTORC1 (p = 0.036), compared to controls; protein levels of mTORC2 (p = 0.927) and β-Catenin (p = 0.936) were unchanged (Figure 7). By assessing the protein levels of the main mTOR signaling pathway, we confirm that developmental Mn exposure causes lasting alterations to the mTOR signaling pathway in the PFC, which provides important insight into the molecular mechanisms of catecholaminergic hypofunctioning and attentional dysfunction in our animal model of childhood developmental Mn exposure and inattention.

**Figure 7:**
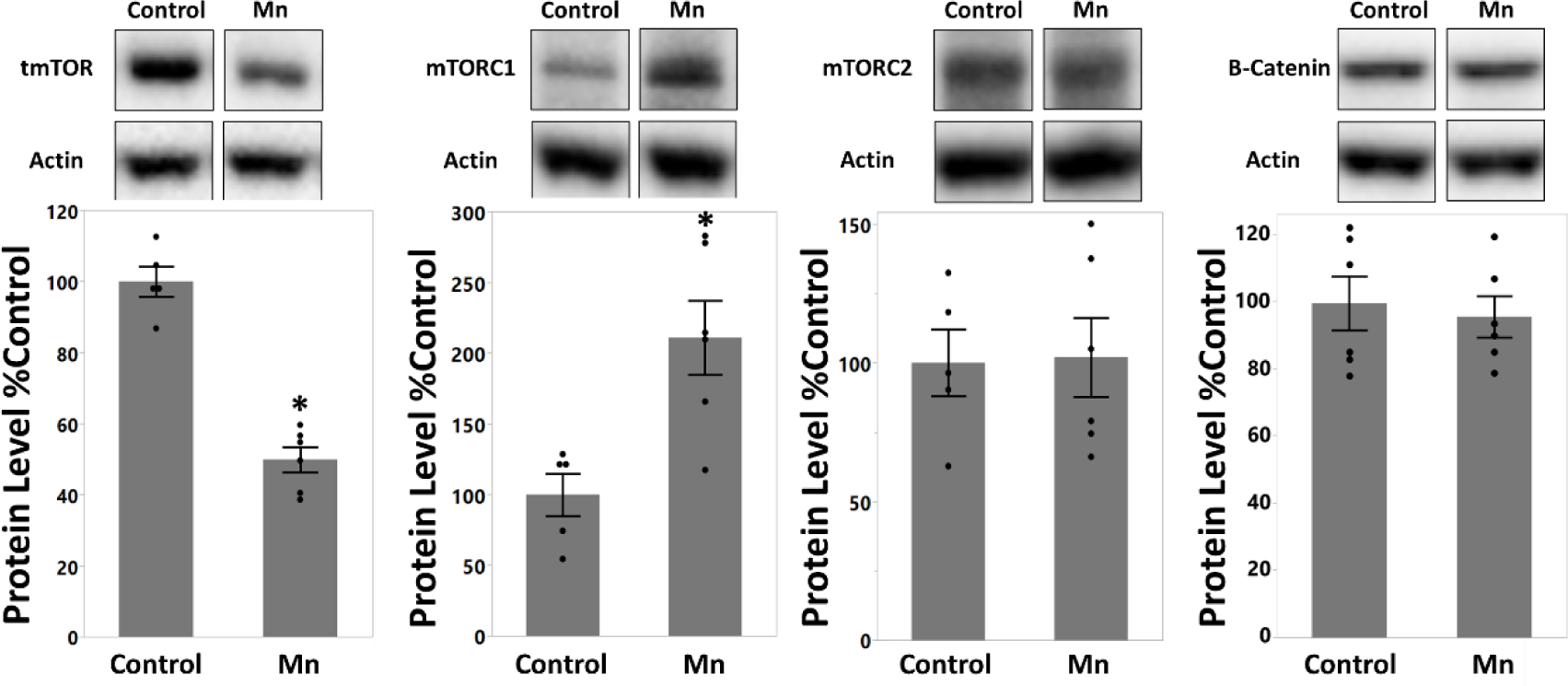
mTOR signaling is established as a novel mechanistic target of Mn neurotoxicity. Protein levels of total mTOR, mTORC1, mTORC2, and β-Catenin in prefrontal cortex of PND 66 rats. Data are presented as normalized to % Control. Mn reduced total mTOR, increased mTORC1, and did not measurably change mTORC2 or β-Catenin protein levels. Statistical comparisons using Wilcoxon: Total mTOR (p = 0.008), mTORC1 (p = 0.036), mTORC2 (p = 0.927), β-Catenin (p = 0.936). Data are means ± SEM, n = 5-6 rats per treatment group.

### 3.8. Developmental Mn exposure increased blood Mn levels and these levels normalize by PND 66

Blood Mn analyses in control and developmentally Mn-exposed rats shows that, following weaning (PND 24) the oral Mn exposure regimen used here increased blood Mn to levels comparable to environmentally-exposed infants, and that by young adulthood (PND 66), blood Mn levels had returned to control levels. Specifically, our developmental Mn exposure regimen over PND 1 – 21 significantly increased blood Mn levels of PND 24 rats to ∼496 ng/mL, compared to 24 ng/mL in controls (p < 0.001) (Supplemental Figure 9). Following cessation of Mn exposure on PND 21, blood Mn levels in the Mn group (15 ng/mL) declined to levels comparable to controls (11 ng/mL) in PND 66 animals. The findings support that the attention deficits and molecular changes caused by developmental Mn exposure are due to lasting neurochemical changes, and cannot be attributed to elevated Mn levels concurrent with behavioral testing and the neuromolecular measurements.

## 4. Discussion

Elevated Mn exposure is associated with inattention, impulsivity, and psychomotor deficits in children, and is recognized as an environmental risk factor for ADHD (2-18, 75). Animal model studies have demonstrated that developmental Mn exposure can cause these ADHD-like attention deficit symptoms, which are associated with lasting hypofunctioning of the catecholaminergic system in the PFC and striatum (22–29). However, the molecular mechanism(s) that underlie these lasting Mn deficits remain poorly understood. To address this, we used a rat model of childhood environmental Mn exposure and lasting attentional dysfunction to investigate whether Mn exposure leads to lasting alterations of PFC epigenomic and transcriptomic domains that may help explain the lasting catecholaminergic and attention deficits reported previously.

Here, we found that developmental Mn exposure causes lasting attentional deficits that are accompanied with reduced expression of the key catecholaminergic system genes tyrosine hydroxylase and dopamine transporter, as well as the epigenetic regulator DNA methyltransferase 3a in the PFC. These findings are consistent with our previous findings showing reduced tyrosine hydroxylase and dopamine transporter protein expression in Mn-exposed animals, while also opening up the possibility of epigenetic regulation mechanisms as an underlying feature of how developmental Mn may cause attention deficits that last into adulthood long after elevated exposure has ended (22). In gaining a broader understanding of how developmental Mn exposure alters the epigenome and transcriptome, our Gene Ontology results showed changes in the broader functional categories of inflammation, epigenetic regulation, cell development, and hypofunctioning neuronal systems. These are important new findings which illuminate potential neural changes responsible for the developmental Mn exposure associated functional impairments.

Upon investigating potential IPA-identified causal and upstream regulators of our transcriptome, our findings further showed specific dysregulation of mTOR and Wnt signaling pathways. Although β-Catenin protein levels did not measurably change with developmental Mn exposure, we did measure altered mTOR pathway protein levels, which supports our hypothesis of altered mTOR signaling as an underlying mechanism for how developmental Mn leads to lasting hypofunctioning of the catecholaminergic system in the PFC. Our findings corroborate the limited published data on Mn and mTOR pathway activation; specifically, one prior rodent study found that low dose developmental Mn exposure increased mTORC1 protein expression (76).

The identification of altered mTOR signaling as a key upstream regulator that can account for the many of the lasting catecholaminergic gene and protein expression changes caused by developmental Mn exposure is a significant finding of our study. mTOR functions as a core component of two distinct protein complexes, mTOR complex 1 (mTORC1) and mTOR complex 2 (mTORC2) (78–79). These two complexes are known to regulate different neurodevelopmental processes, such as neuron size, dendritic growth and maturation, and axon elongation (80–81).

Our differential methylation, differential gene expression, IPA integration, and protein level findings provide evidence that mTORC1 is an activated IPA causal regulator of our Mn-exposure transcriptomic phenotype that also has increased protein expression. Prior studies have demonstrated that persistently elevated mTORC1 signaling blocks canonical D1R signaling, which has been shown to alter dopamine signaling pathway gene expression, cortico-striatal plasticity, and cognitive functions in mice (54–58). Furthermore, DA-Tsc1-KO mice that have a loss of tuberous sclerosis complex 1 (*Tsc1*) function, an upstream regulator of mTOR activity, leads to constitutive activation of mTORC1, striatal dopamine neuron somatodendritic hypertrophy, reduced intrinsic excitability, altered axon terminal structure, and reduced dopamine release (82). Interestingly, these mice have increased tyrosine hydroxylase synthesis, but reduced dopamine release due to decreased vesicle function; these changes are associated with reduced cognitive flexibility outcomes, which were rescued by genetic reduction in activated mTORC1 signaling (82).

Although we were unable to detect a significant reduction in mTORC2 protein levels, our differential methylation, gene expression, and Ingenuity Pathway Analysis indicates inhibited mTORC2 signaling as a consequence of developmental Mn exposure. This predicted mTORC2 dysregulation is evident through Rictor, an essential component of the mTORC2 complex, measured here as hypermethylated and IPA predicted inhibited. Our findings of *Wnt*, *Gsk3b*, *Akt,* and *Tsc1-Tsc2* complex differential methylation and IPA dysregulation as well as *Tsc2* differential gene expression give evidence of dysregulated mTORC2 based on their respective pathway interactions (Figure 8) (89–92). Mice without functioning *Rictor* (nRictor KO) have demonstrated novelty-induced hyperactivity, decreased dopamine bioavailability, and elevated dopamine receptor D2 protein expression (83). These findings specifically mirror our previous findings that developmental Mn exposure significantly decreases dopamine and norepinephrine evoked release, and that developmental Mn exposure increases dopamine receptor D2 protein expression levels in the prefrontal cortex by about 200% (22, 29). Interestingly, based on previous research (93–94), elevated dopamine receptor D2 protein expression and/or the inhibition of *Akt* caused by inhibited mTORC2 can contribute to our ∼80% Mn-induced decrease in dopamine transporter gene expression (present study) and a similarly decreased protein expression findings shown in our prior studies (22).

**Figure 8:**
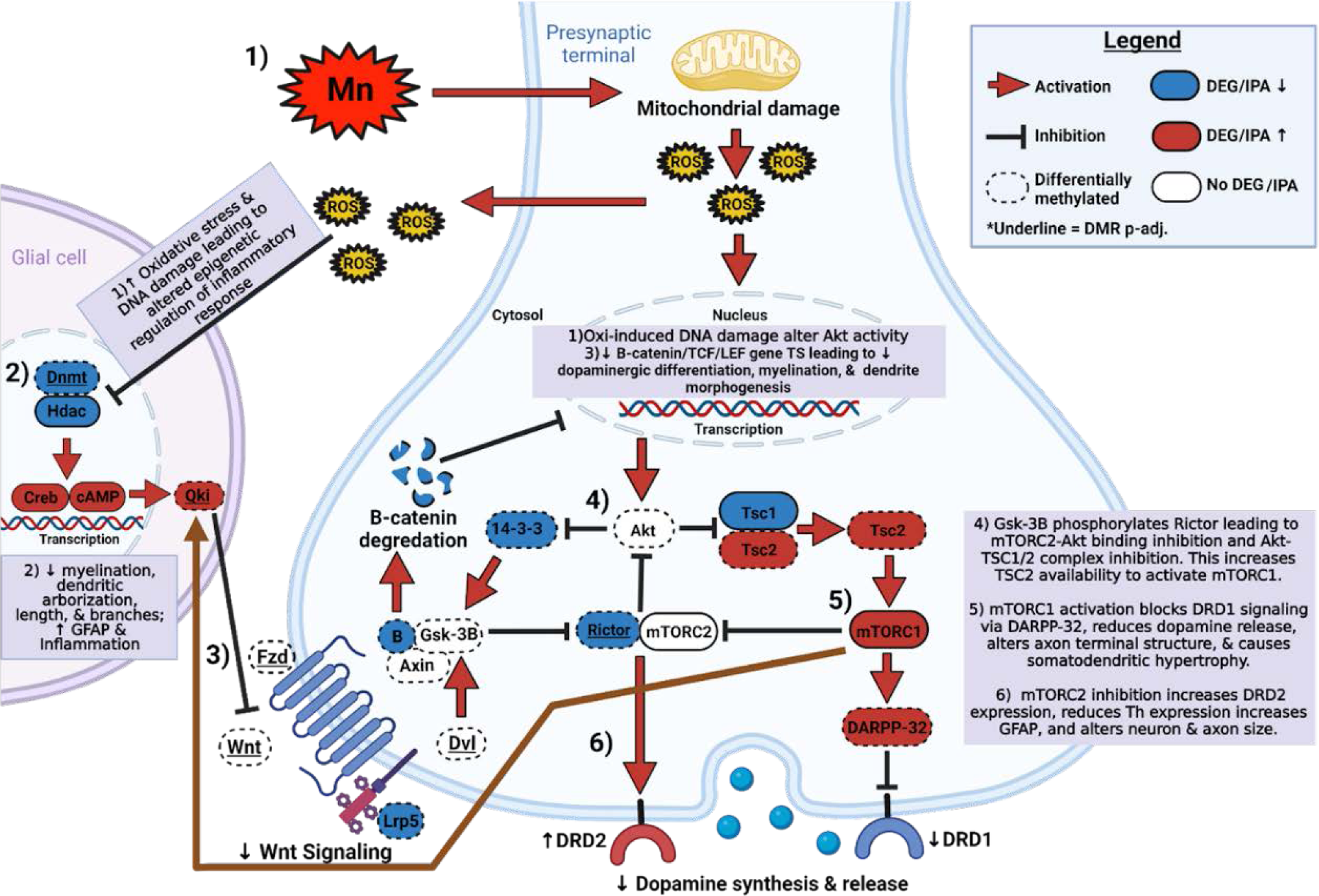
Proposed mechanism of Mn neurotoxicity based on integration of p-unadjusted DMRs, DEGs, IPA upstream and casual regulators, and peer reviewed scientific literature. Briefly, Mn exposure may induce oxidative stress and DNA damage leading to altered epigenetic regulation of inflammatory response as well as direct change to Akt activity. This altered regulation leads to reduced Wnt signaling and as a result increased mTORC1 and decreased mTORC2 regulation. Altered mTORC1/mTORC2 impair the catecholaminergic system and perpetuates a continued inflammatory environment reflecting our Mn phenotype of lasting catecholaminergic system hypofunctioning and attention deficits. Legend: Red arrow is activation; Black Line with bar is inhibition; Dashed circle is differentially methylated; Blue Fill is DEG/IPA downregulated prediction; Red Fill is DEG/IPA upregulated prediction; No fill is no DEG/IPA evidence; Underlined name is DMR p-unadjusted and p-adjusted significance. Figure was made in Biorender Premium.

We propose a series of mechanisms that may help explain how developmental Mn exposure leads to lasting deficits in attentional function in our rat model (Figure 1 and Figure 8). This model integrates our new understanding of how Mn exposure affects differentially methylated and expressed genes, including causal and upstream regulator genes. Briefly, we propose that developmental Mn exposure induces oxidative stress and DNA damage leading to altered epigenetic regulation of inflammatory responses, as well as direct change to Akt activity (Figure 8, Step 1-3) (30-32, 84-90). This leads to reduced Wnt signaling, and as a result increased mTORC1 and decreased mTORC2 regulation (Figure 8, Step 4-6) (56, 91–94). Altered mTORC1/mTORC2 signaling impairs the catecholaminergic system and promotes a proinflammatory environment of activated glial cells, resulting in a hypofunctioning catecholaminergic system and attention deficits (Figure 8, Step 4-6) (22-29, 54-58, 95-98). This activation of glial cells is consistent with our prior findings that developmental Mn exposure leads to lasting rat PFC GFAP and pro-inflammatory astrocyte activation (22).

In summary, our findings demonstrate that developmental Mn exposure causes lasting differential methylation and gene expression changes in inflammatory, epigenetic regulation, cell development, and hypofunctioning neuronal systems with reduced expression of the catecholaminergic system genes tyrosine hydroxylase and dopamine transporter, as well as DNA methyltransferase 3a, in the prefrontal cortex. Additionally, developmental Mn exposure causes lasting methylation, gene expression, and protein level dysregulation in mTOR and Wnt signaling pathways and these resulting dysregulations can induce lasting prefrontal cortex catecholaminergic system hypofunctioning, and may underlie or contribute to the lasting deficits in attention and impulse control reported here and in our prior studies (22, 27–29). Importantly, we establish early development as a critical period of susceptibility to lasting deficits in attentional function caused by elevated environmental toxicant exposure. Our findings deepen our understanding of the lasting epigenetic and gene transcriptomic changes caused by developmental Mn exposure, and how these changes may lead to lasting catecholaminergic system dysfunction. Specifically, our findings of altered mTOR pathway regulation may aid potential therapeutic studies on the environmental exposure contribution to neurological disease.

## 5 Acknowledgements

We thank Ellie Fisher, Samantha Gorman, Belinda Huang, Marissa Mena, Angelica Montevirgen, Naomi Schlipp, Garrick Sin, Sheila Sharma, Shannon Twardy, and David Woodfin for assistance with the attention series testing and Tom Jursa for tissue analyses for Mn concentrations. We also would like to thank the UC Davis Genome Core conducting the gene expression and methylation sequencing as well as the UCLA Technology Center for Genomics & Bioinformatics for sending us the initial Ingenuity Pathway Analysis data for our analysis. Funding was from the National Institutes of Environmental Health Sciences (#R01ES028369).

