## Supplemental materials for "Developmental Manganese Exposure Causes Lasting Attention Deficits Accompanied by Dysregulation of mTOR Signaling and Catecholaminergic Gene Expression in Brain Prefrontal Cortex"

### Index

*Supplemental Methods*……………………………………………..pg. 2 - 7

*Supplemental Tables*……………………………………………..pg. 8

*Supplemental Figures*…………………………………………..…..pg. 9 - 17

*Supplemental References*……………………………………..…pg. 18 - 20

**<u>Supplemental Methods:</u> (Title # Corresponds to Main Manuscript)**

#### 2.2. Manganese Dosing Regimen and Experimental Design

Neonatal rats were orally exposed to a Mn dose of 50 mg Mn/kg/day starting on PND 1 through weaning on PND 21 (early postnatal Mn exposure; n=64 per treatment group). The early window of postnatal Mn exposure in rats corresponds to important frontal-cortical-striatal developmental events, including formation of DA and NE projections to the PFC, that are comparable to the gestational third trimester through adolescence in humans (1-4). For Mn dosing, a 225 mg Mn/mL stock solution of MnCl2 was prepared by dissolving MnCl2·4H2O with Milli-Q™ water; aliquots of the stock solution were diluted with a 2.5% (wt/vol) solution of the natural sweetener stevia to facilitate oral dosing of the pups. The stock solutions were made fresh weekly. Control rats received the vehicle solution. This Mn exposure regimen is relevant to children exposed to elevated Mn via drinking water, diet, or both; pre-weaning exposure to 50 mg Mn/kg/day produces a relative increase in Mn intake that approximates the increase reported in infants and young children exposed to Mn-contaminated water or soy-based formulas (6-13). Rats were allowed to age into adulthood and behavioral analysis was conducted at PND 73-102.

#### 2.3. Focused Attention

To assess focused attention the 5-Choice Serial Reaction Time Task (5-CSRTT) was used as previously described (9-10). Briefly, developmentally exposed rats began testing at about PND 45, with food magazine and nose-poke training for 1 week followed by two 5-choice visual discrimination tasks (1: fixed cue duration of 15 seconds/ 2: fixed cue duration of 1 second). Once the animal attains 80% correct on two of three successive sessions in the visual discrimination tasks, the animal moves on to the Focused Attention Tasks. Both focused attention tasks assessed the ability of the rats to detect and respond to a brief visual cue presented unpredictably in time and location (one of the five response ports). Two focused attention tasks (#1 and #2) were administered over PND 73–86, and PND 88–102, respectively. The Focused Attention Task 1 used variable pre-cue delays of 0, 1, 2, or 3 (on top of the standard 3 s delay to allow for animal reorientation) and a fixed visual cue duration of 0.700 sec was administered for 12 sessions. In the Focused Attention Task 2 the same variable pre-cue delay of 0, 3, 4, or 5 sec is imposed between trial onset and the presentation of the visual cue, but the duration of the light cue is either 0.400 or 0.700 sec which presents an additional attentional challenge for the rat. The Focused Attention Task 2 was administered for 12 sessions (150 trials/session), the time required for most rats to achieve stable performance. The variable pre-cue delays are presented pseudo-randomly with the provision that each delay is used equally in each session. All rats were weighed and tested 6 days/week throughout training and testing. Behavioral assessment occurred during the active (dark) period of the diurnal cycle at the same time each day and in the same chamber for each individual rat. All behavioral testing was conducted by individuals blind to the treatment condition of the subjects. All rats were maintained on a food restriction schedule with water available ad lib throughout behavioral assessment, as described previously (9-10).

### 2.4. Targeted Catecholamine and Epigenetic System Gene Expression Analysis

In order to determine gene expression of key catecholaminergic genes (Th, Dat, Drd2) and epigenetic modulators (Dnmt1, Dnmt3a, Dnmt3b, Hdac3, and Hdac4) we conducted RT-qPCR on PFC tissue from PND 66 control and developmental Mn exposed rats (6-7 rats/treatment). Briefly, fresh-frozen tissue punches (25-30 mg) were taken from the anterior cingulate cortex/prefrontal cortex area (PFC) (Paxinos and Watson Rat Brain Atlas: 2.16-0.77 mm anterior to bregma) over dry-ice and underwent Dounce Homogenization and total RNA extraction using the Quick-DNA/RNA™ Miniprep Kit (Zymo: Cat. #d7001) following the manufacturer's instructions. RNA quality and quantity was measured through Nanodrop and RNA band integrity was confirmed through gel electrophoresis prior to DNAse Treatment and Reverse Transcription (RT) using the Invitrogen™ SuperScript™ IV VILO™ Master Mix with ezDNase™ Enzyme kit (Invitrogen™: Cat. #11766050) following the manufacturer's instructions. RT-qPCR cycle quantity thresholds were measured on 1 - 5 ng cDNA by the Bio-Rad CFX96 using the ThermoFisher Scientific TaqMan Advanced Master Mix (Applied Biosystems: Cat. #4444556) and TaqMan primers (Supplemental Table 1) following the manufacturer's protocol. Three reference genes were assessed (Gapdh, ActB, and Ubc) and the tool Normfinder confirmed our most stable reference gene combination for gene expression analysis was to normalize each target gene to the average RT geometric mean of Gapdh, ActB, and Ubc cycle quantity threshold (14-15). NormFinder determines the stability of the candidate reference genes by measuring the intra- and intergroup variation between user specified groups (e.g., treated and control groups) first. Stability values for each candidate gene are then calculated by adding the two sources of variation. The lowest stability value means the most stable expression (15).

RT-qPCR was performed with two RT reaction replicates per animal (n=6/7), and 3 sample replicates per RT for each animal based on best recommended RT-qPCR practices to reduce experimental variability (16). The standard $\Delta$Ct method of gene expression quantification was used where the $\Delta$Ct method compares relative expression of pairs of candidate genes within each sample in order to determine whether candidate genes are differentially expressed by a treatment condition. If the $\Delta$Ct value of the two genes fluctuates when analyzed in different treatment conditions, it means that one or both genes are variably expressed (17). RT-qPCR data statistical analyses were done on $\Delta$Ct values by non-parametric Wilcoxon signed-rank test analysis and $p \leq 0.05$ deemed significance.

### 2.5. Differential DNA Methylation and Functional Pathway Analysis

In order to determine whether developmental Mn exposure led to lasting alterations of genome methylation status that may contribute to a Mn-induced hypofunctioning catecholaminergic system, we conducted reduced representation bisulfite sequencing (RRBS). DNA was extracted from the same PND 66 fresh-frozen PFC samples detailed in Section 2.4. using the Quick-DNA/RNA™ Miniprep Kit (Zymo: Cat. #d7001) following the manufacturer's instructions. 500 ng of DNA was bioanalyzed for DNA integrity, bisulfite converted for RRBS using the Premium RRBS kit (Diagenode: Cat. #C02030033), and sequenced by Illumina HiSeq PE150 for about 40 million reads each sample by the University of California Davis Genome Center (n= 3/treatment).

To obtain differentially methylated regions (DMRs) between control and Mn groups Galaxy Pipeline Analysis was conducted. Briefly, FastQ sequence files underwent automatic quality and adapter trimming using TrimGalore with a <30 Phred quality score. FastQC/MultiQC were then used to assure the trimmed reads passed the default algorithm's quality parameters and then BWA-meth was used to align the reads to the rat genome (rn6) reference sequence. Methyldackel was then used to assess methylation bias and to extract methylation values from the sequences, and then Wig/BedGraph-to-bigWig was used to visualize the methylation levels. The extracted methylation values between control and Mn exposed rats were then assessed using Metilene, which gave an output of all significant DMRs ($p < 0.05$) (18-21). These significant DMRs (unadjusted and adjusted $p < 0.05$) were annotated by chromosome to the rat (rn6) genome using the UCSC Genome Integrator, giving an output of the DMR associated gene by chromosome location. These DMR chromosome locations were then visualized and manually confirmed using the Integrative Genomics Viewer (IGV) sliding window (22). By aligning the DMRs to the rat genome we are able to associate the DMRs with specific genes as well as determine the location of DNA methylation within the gene, such as the promoter region or a specific exon; the promotor region of a gene was determined as between 1,500 bp upstream and 500 bp downstream of transcription start sites (23). These identified genes are referred here as differentially methylated (DM).

Next, fold-change values were calculated for each DM gene as the difference of the mean methylation value of the control versus Mn PFC samples. All 4,250 $p<0.05$ significant genes were then ranked using the $\text{Log}_{10}$ of their respective p-value multiplied by their calculated fold-change, and then Fast Preranked Gene Set Enrichment Analysis (FGSEA) was performed to determine the associated functions of the DM gene products through Gene Ontology (GO) Biological Processes (BP) (24). GO BP represents a specific objective that the organism is genetically "programmed" to achieve, which is often portrayed as an outcome or ending state (25-26). Results are provided in hierarchical terms (parent and child terms) that helps identify upstream and downstream processes and establish similarity connections. GO BP groups were further reduced, filtered, and visualized by the tool REVIGO's algorithm for semantic similarity of parent and child terms (27). Additionally, Kyoto Encyclopedia of Genes and Genomes (KEGG) analysis were conducted on all significant 4250 DMR associated genes ($p < 0.05$ unadjusted) using EnrichR (28). KEGG analysis integrates 16 databases that allows us to further understand gene product functions by putting them in the context of disease and pharmaceutical outcomes (29).

### 2.6. Differential Expressed Genes and Functional Pathway Analysis

To determine whether developmental Mn exposure was associated with altered gene expression 1 ug total RNA aliquots from the same PND 66 PFC RNA extraction in Method 2.4 were analyzed for Bioanalyzer quality assessment, library preparation, and 3' Tag RNA-sequencing by the University of California Davis Genome Center. 3' Tag RNA-seq generates a single initial library molecule per transcript, complementary to 3′ end sequences, in contrast to traditional RNA-seq which generates sequencing libraries for the whole transcript. This method 3′ generates low-cost and low-noise gene expression profiling data suitable for differential gene expression studies (30-32). Subsequently, all FastQ files were processed using established Geneious Prime RNA-sequencing tools (Geneious Prime 2022.0.). Briefly, TrimGalore was used to remove adapters and trim reads for a <30 Phred Quality Score. Geneious Prime Genome Alignment was then used, to align the reads to the rat (rn6) genome reference sequence, followed by Geneious Prime Annotation analyses to annotate and calculate individual animal gene expression values. DESeq2 was then used to calculate differentially expressed gene (DEG) values between control and developmental Mn exposed PFC brain samples (33).

Next, all DEGs were ranked using the $Log_{10}$ of their respective p-value multiplied by their calculated fold-change and Fast Preranked Gene Set Enrichment Analysis (FGSEA) was then used to determine the DEG's Gene Ontology (GO) associated Biological Processes (BP), Molecular Functions (MF), and Cellular Components (CC) subtypes. The relationship between these GO analysis subtypes is defined by genes encoding gene products in which then carry out molecular-level functions (MF) within specific cellular locations (CC), and these molecular processes contribute to a larger biological objective (BP) (25-26). GO BP groups were further reduced, filtered, and visualized using REVIGO's algorithm for semantic similarity of parent and child terms (27). These reduced GO groups then were categorized into four functional categories (inflammation, epigenetics, cell development, and neuronal function), based on their REVIGO similarity results, Rat Genome Data base functions, and *a priori* understanding of Mn mechanisms of neurotoxicity from the literature. Their respective leading edge contributing genes (core genes of high scoring gene sets that contribute to the enrichment score) and fold-changes are displayed in a heat map generated by the tool Clustergrammer (34). Additionally, KEGG analysis was conducted on all significant DMR associated genes ($p < 0.05$ unadjusted) using EnrichR (28).

### 2.7. Differential Methylation and Differential Expression Integration

In order to determine the Gene Ontology biological processes associated with both differential methylation and differential gene expression of our Mn neurotoxicity attention deficit phenotype, we conducted an integrated analysis of DMR and DEG data using the Genomics Tools Venn-Diagram web program to compare and visualize the upregulated and downregulated Gene Ontology biological processes group lists shared by both DMR associated genes and DEGs using REVIGO, and the four *a priori* categories of biological function (inflammation, epigenetics, cell development, and neuronal function) described in Methods 2.5. Genomics Tools Venn-Diagram web program comparison was performed differentially methylated and expressed genes to further determine genes that were both differentially methylated and differentially expressed. Clustergrammer was then used to visually display DMR and DEG expression levels and the gene region location of the DMRs (e.g., promoter, exon, intron, etc.) were determined using the UCSC Genome Integrator and IGV slide sorting noted in Methods 2.5.

In order to determine which genes may be regulators of our lasting Mn-induced gene expression molecular phenotype, we used Ingenuity Pathway Analysis (IPA) by UCLA Technology Center for Genomics & Bioinformatics on all p-unadjusted for multiple comparison genes conducted from DESeq2 to identify significant causal and upstream regulator genes IPA Upstream Regulator Gene analysis can identify the cascade of upstream transcriptional regulators that may explain the observed gene expression changes in a database (35). IPA Causal Gene Analysis expands on upstream regulator analysis by including gene regulators that are known to regulate the identified upstream regulators of the data set (35). Genes that were identified as upstream and causally-related to DEGs were then integrated with differentially methylated and expressed genes through Genomics Tools Venn-Diagram web program comparison to identify genes that may be mechanistically associated with the Mn-neurotoxicity DEG phenotype. Finally, in order to further strengthen our assessment of the significance of identified causal and upstream regulators we compared the DMR and DEG lists to the most significant activation value regulator

Search Tool for the Retrieval of Interacting Genes/Proteins (STRING) gene interaction pathway lists (36).

#### 2.8. Total and Phosphorylated mTOR Protein Levels

To confirm mTOR pathway proteomic dysfunction Western blots were conducted for total mTOR, mTORC1, and mTORC2 protein levels. Briefly, protein was extracted from PFC brain punches of PND 66 control and Mn rats. Tissue punches (list punch diameter, yielding 25-30 mg) were taken from 1.39 mm thick brain sections (2.16-0.77 mm anterior to bregma-Paxinos and Watson's The Rat Brain in Stereotaxic Coordinates. 7th Edition.), homogenized on ice using a Dounce homogenizer, and brief sonicated in RIPA buffer (Millipore-Sigma: Cat. #20-188) and protease inhibitor cocktail (ThermoFisher: Cat. #PI78441). Samples were vortexed and centrifuged at 11,000 × g for 10 minutes at 4°C and the supernatant was collected. Lysate protein concentrations were determined by BCA following the manufacturer’s protocol (Fisher Scientific: Cat. #23227). Protein lysate aliquots were prepared for Western blot analyses by adding standard NuPAGE LDS Buffer (ThermoFisher: Cat. #NP0008), reducing agents (Life Technologies: Cat. #NP0004), and Milli-Q water, mixed, and heated to 70°C for 10 minutes, then stored until at -20°C until use. Western blots were run on the Invitrogen XCell Sure Lock system using NUPAGE MOPS SDS Running Buffer (Life Technologies: Cat. #NP0001) at constant 100 V c for 2.5 hours at 4°C with 20 µg of protein per lane and electrophoresed on 4-12% NuPAGE 15 well gels (Life Technologies: Cat. #NP0323BOX). Protein was transferred to PVDF membranes using the Invitrogen XCell II Blot Module system overnight at 4°C; success of protein transfer was determined through standard Ponceau (PVDF membrane) and Coomassie staining (gel). PVDF membranes were blocked in 5% non-fat milk tris buffer saline/Tween-20 for 2 hours at room temperature (RT) on a shaker, and then incubated in primary antibody (1:1000) overnight on a shaker at 4°C (for antibodies see Supplemental Table 2). Following overnight primary antibody incubation, membranes were washed 6x for 5 min each, blocked in 5% normal donkey serum/TBST for 1 hour RT, incubated in donkey anti-mouse IgG-HRP secondary antibody (1:10000; ThermoFisher: Cat. #A16017), washed 6x for 5 min each, and visualized using SuperSignal™ West Pico PLUS Chemiluminescent Substrate (Thermofisher: Cat. #34578) on a Bio-Rad ChemiDoc instrument.

Western blot band densitometry was quantified using Bio-Rad Image Lab volume quantification software. Briefly, lanes and bands were identified on each membrane image and the volume tools were used to measure each target protein band. Then each target protein value was normalized to a background no signal region and then to each respective beta-actin band value for each sample. 6 control and 6 Mn rats, run in consecutive order with 2 gels/target protein were analyzed with Wilcoxon statistical analysis to establish $p < 0.05$ statistical significance.

#### 2.9. Mn Blood Levels

Blood Mn concentrations were measured in the same PND 66 rats used in the reported molecular outcomes and were measured as previously published (6-13). Briefly, rats underwent euthanasia via $CO_2$ inhalation, whole blood (2 – 3 mL) was collected from the left ventricle of the surgically-exposed heart put into EDTA vacutainers on dry ice and then stored at −20 °C for later analyses. Aliquots of whole blood were digested overnight at room temperature with 16 M HNO3 (Optima grade, Fisher Scientific), followed by addition of $H_2O_2$ and Milli-Q water. Digestates were centrifuged (13,000 x g for 15 min.) and the supernatant collected for Mn analysis. Rhodium was added to sample aliquots as an internal standard. Mn levels were determined using a Thermo Element XR inductively coupled plasma–mass spectrometer, measuring masses 55Mn and 103Rh (the latter for internal standardization). External standardization for Mn used certified SPEX standards (Spex Industries, Inc., Edison, NJ). National Institutes of Standards and Technology SRM 1577b (bovine liver) was used to evaluate procedural accuracy. The analytical detection limit for Mn in blood was 0.04 ng/mL, respectively.

**Supplemental Tables:**

| Gene | Symbol | ThermoFisher Assay Id | Amplicon size |
|---|---|---|---|
| Tyrosine hydroxylase | Th | Rn00562500_m1 | 60 |
| Solute carrier family 6 member 3 (dopamine transporter) | Dat (Slc6a3) | Rn00562224_m1 | 111 |
| Dopamine receptor D2 | Drd2 | Rn00561126_m1 | 64 |
| DNA methyltransferase 1 | Dnmt1 | Rn00709664_m1 | 62 |
| DNA methyltransferase 3a | Dnmt3a | Rn01027162_g1 | 59 |
| DNA methyltransferase 3b | Dnmt3b | Rn01536419_m1 | 54 |
| Histone deacetylase 3 | Hdac3 | Rn00584926_m1 | 58 |
| Histone deacetylase 4 | Hdac4 | Rn01427040_m1 | 60 |
| Glyceraldehyde 3-phosphate dehydrogenase | Gapdh | Rn99999916_s1 | 87 |
| Beta-actin | Actb | Rn00667869_m1 | 91 |
| Ubiquitin C | Ubc | Rn01789812_g1 | 88 |

**Supplemental Table 1:** Each gene expression assay consists of a mixture of sequence specific forward/reverse primers and a FAM-labeled probe. Primer/probe sets that are labeled "m1" cross an exon-exon boundary, labeled "g1" cross an exon-exon boundary but may still detect genomic DNA, and those labeled "s1" are found within a single exon (i.e. detects genomic DNA).

| Protein | Vendor | Catalog Number | kDa |
|---|---|---|---|
| Total mTOR | Santa Cruz Biotech. | #sc-517464 | 245 |
| Phospho-mTORC1 | Santa Cruz Biotech. | #sc-293133 | 220 |
| Phospho-mTORC2 | Santa Cruz Biotech. | #sc-293132 | 220 |
| Beta-catenin | Santa Cruz Biotech. | #sc-7963 | 90 |
| Beta-actin | Santa Cruz Biotech. | #sc-47778 | 42 |

**Supplemental Table 2:** Primary antibodies used for Western blot analysis of total mTOR, phospho-mTORC1, phosphor-mTORC2, and reference protein beta-actin.

### Supplemental Figures:

#### Top 10 Significant KEGG 2021 Terms

##### Hypermethylated

| Term | P-value | Overlap Genes |
|---|---|---|
| Hippo signaling pathway | 4.638057e-14 | GSK3B, PATJ, WNT2B, FZD10, ACTB, ACTG1, GLI2, SOX2, PPP1CC, CCND1, YWHAQ, CDH1, MYC, CCN2, TEAD2, TEAD3, YWHAH, APC2, TEAD4, PRKCI, CSNK1E, AXIN2, WNT9A, TGFBR2, LATS1, LATS2, AJUBA, LLGL1, LLGL2, CRB2, TCF7, LEF1, PARD6G, WNT8B, PRKCZ, NKD1, SAV1, NKD2, PARD6B, DVL1, DVL2, TP53BP2, DVL3, CTNNA2, WNT1, WNT3, WNT4, FZD1, SMAD1, WNT10B, WWTR1, TCF7L1, FZD2, SMAD3, TGFB1, FZD5, FZD4, FZD7, WNT7B, FZD9, BMP8B, WNT7A, BMP7, SMAD7, MOB1B, DLG4, CTNNB1, DCHS1, FAT4, BMPR1A |
| Gastric cancer | 6.756033e-12 | GSK3B, CDKN1A, CDKN1B, WNT2B, FZD10, FGF3, FGF5, SHH, CCND1, FGF9, CDH1, MYC, AKT1, HRAS, APC2, MAP2K2, AXIN2, WNT9A, TGFBR2, CCNE2, RAF1, SHC4, SHC2, TERC, TCF7, LEF1, LRP5, WNT8B, PIK3R2, RXRB, TERT, DVL1, DVL2, DVL3, MAPK1, CTNNA2, WNT1, WNT3, WNT4, MAPK3, FZD1, WNT10B, CDKN2B, TCF7L1, FZD2, SMAD3, TGFB1, FZD5, GADD45B, CDX2, FZD4, GADD45A, FZD7, WNT7B, FZD9, WNT7A, MLH1, FGF18, RPS6KB2, CTNNB1, GRB2, KRAS |
| Proteoglycans in cancer | 1.031577e-11 | CDKN1A, WNT2B, ITGB5, ITGB3, IHH, FZD10, ACTB, ACTG1, PPP1CC, SHH, CCND1, MYC, AKT1, TIMP3, ITGAV, RAC1, HRAS, PRKCG, VAV3, MAP2K2, ARHGEF12, PDPK1, WNT9A, NUDT16L1, ANK1, TIAM1, SMO, ITGA5, RAF1, PPP1R12C, HBEGF, TLR2, CAMK2B, SDC4, ROCK1, SRC, SDC2, PXN, WNT8B, PIK3R2, ITPR3, HOXD10, SLC9A1, ERBB3, ERBB4, GPC1, DROSHA, MAPK1, PLCG1, WNT1, CAMK2G, WNT3, EIF4B, WNT4, MAPK3, FZD1, WNT10B, FZD2, TGFB1, FZD5, FZD4, FZD7, WNT7B, PTCH1, FZD9, WNT7A, PTPN11, VEGFA, MAPK13, MAPK11, TFAP4, RPS6KB2, MDM2, SDC1, CTNNB1, GRB2, KRAS |
| Basal cell carcinoma | 1.399522e-11 | GSK3B, CDKN1A, WNT2B, LEF1, TCF7, WNT8B, FZD10, GLI1, GLI3, GLI2, SHH, DVL1, DVL2, DVL3, WNT1, WNT3, WNT4, APC2, FZD1, WNT10B, TCF7L1, FZD2, FZD5, GADD45B, FZD4, GADD45A, FZD7, PTCH1, WNT7B, FZD9, WNT7A, AXIN2, WNT9A, SMO, CTNNB1 |
| Signaling pathways regulating pluripotency of stem cells | 3.284718e-11 | GSK3B, WNT2B, FZD10, SOX2, MYC, AKT1, JAK2, HRAS, JAK3, APC2, MAP2K2, PAX6, AXIN2, WNT9A, DUSP9, IL6ST, RAF1, HOXD1, TCF7, WNT8B, PIK3R2, ACVR1B, DVL1, DVL2, DVL3, MAPK1, WNT1, WNT3, WNT4, MAPK3, FZD1, SMAD1, WNT10B, PCGF6, FZD2, SMAD3, FZD5, SETDB1, FZD4, PCGF5, FZD7, PCGF2, WNT7B, FZD9, LIF, WNT7A, INHBB, KLF4, TBX3, MAPK13, MAPK11, ID3, CTNNB1, GRB2, LHX5, KRAS, TCF3, FGFR3, BMPR1A |
| Cushing syndrome | 5.012930e-11 | GSK3B, CDKN1A, CDKN1B, WNT2B, ASH2L, AHR, FZD10, CCND1, CREB3L1, APC2, USP8, MAP2K2, AXIN2, WNT9A, NR5A1, ARMC5, CCNE2, ATF4, CREB5, CAMK2B, KMT2A, ATF6B, TCF7, LEF1, WNT8B, ITPR3, ADCY2, CACNA1C, ADCY8, CACNA1H, CRHR1, ADCY6, CACNA1G, ADCY5, RASD1, GNA11, DVL1, DVL2, DVL3, MAPK1, WNT1, CAMK2G, WNT3, WNT4, MAPK3, FZD1, WNT10B, CDKN2B, TCF7L1, FZD2, FZD5, FZD4, FZD7, WNT7B, FZD9, WNT7A, CDK6, GNAQ, GNAS, CTNNB1, KCNK2, KCNK3 |
| Pathways in cancer | 2.279027e-10 | KEAP1, FZD10, FGF3, FGF5, EDNRA, FGF9, CCND1, CDH1, MYC, PIM1, AKT1, PRKCG, MAP2K2, IL4R, DAPK2, DAPK3, RUNX1, SMO, CCNE2, COL4A1, IL6ST, RAF1, IFNAR1, NOTCH3, TERC, EPAS1, TCF7, PDGFA, LPAR2, TGFA, PIK3R2, RASGRP2, CSF2RA, FOXO1, PLD2, BCL2L11, TERT, DVL1, DVL2, DVL3, PLCG1, FADD, WNT1, RALGDS, WNT3, WNT4, FZD1, JAG2, STAT5A, WNT10B, JUN, FZD2, SMAD3, TGFB1, FZD5, GADD45B, FZD4, GADD45A, FZD7, ZBTB16, PTCH1, FZD9, NFKB2, CDK6, GNAQ, FGF18, MDM2, GNAS, CYCS, GRB2, FGFR3, NFE2L2, GSK3B, CDKN1A, CDKN1B, HSP90AB1, WNT2B, FLT3, FLT4, LAMC1, GLI1, GLI3, GLI2, IKBKB, CASP9, SHH, CASP7, ITGAV, RAC1, JAK2, JAK3, HRAS, APC2, ARHGEF11, ARHGEF12, APAF1, GSTO1, IFNGR1, ITGA3, PLEKHG5, ... |
| Hepatocellular carcinoma | 2.823376e-10 | GSK3B, CDKN1A, WNT2B, KEAP1, FZD10, ACTB, ACTG1, CCND1, MYC, AKT1, HRAS, PRKCG, APC2, MAP2K2, SMARCC2, GSTO1, AXIN2, WNT9A, TGFBR2, RAF1, SHC4, SHC2, SMARCD2, TERC, TCF7, LEF1, LRP5, TGFA, WNT8B, PIK3R2, TERT, DVL1, DVL2, DVL3, MAPK1, PLCG1, WNT1, BRD7, WNT3, WNT4, MAPK3, FZD1, WNT10B, TCF7L1, FZD2, SMAD3, TGFB1, FZD5, GADD45B, FZD4, GADD45A, BAD, FZD7, TXNRD1, WNT7B, FZD9, WNT7A, SMARCA2, CDK6, RPS6KB2, CTNNB1, GRB2, KRAS, NFE2L2 |
| mTOR signaling pathway | 3.137563e-09 | GSK3B, WNT2B, IRS1, FZD10, IKBKB, DEPTOR, RPS6KA1, MLST8, AKT1, HRAS, PRKCG, SEC13, ATP6V1G2, MAP2K2, MIOS, PDPK1, STRADA, TSC2, WNT9A, RRAGC, AKT1S1, TBC1D7, RAF1, CAB39, NPRL3, LRP5, WNT8B, PIK3R2, WDR24, DVL1, DVL2, DVL3, MAPK1, RICTOR, WNT1, FNIP1, EIF4E, ATP6V1C1, WNT3, EIF4B, ATP6V1C2, WNT4, MAPK3, FZD1, WNT10B, FZD2, WDR59, FZD5, FZD4, FZD7, WNT7B, FZD9, WNT7A, CASTOR1, RPS6KB2, GRB2, KRAS, LAMTOR4 |
| Breast cancer | 3.588958e-09 | GSK3B, CDKN1A, WNT2B, FLT4, FZD10, FGF3, FGF5, CCND1, FGF9, MYC, AKT1, HRAS, APC2, MAP2K2, AXIN2, WNT9A, RAF1, SHC4, NOTCH3, SHC2, TCF7, LEF1, LRP5, WNT8B, PIK3R2, DLL1, DVL1, DVL2, DVL3, MAPK1, WNT1, WNT3, WNT4, MAPK3, FZD1, JAG2, WNT10B, JUN, TCF7L1, FZD2, FZD5, GADD45B, FZD4, GADD45A, FZD7, WNT7B, FZD9, WNT7A, NFKB2, HEYL, CDK6, FGF18, RPS6KB2, CTNNB1, GRB2, KRAS |

##### Hypomethylated

| Term | P-value | Overlap Genes |
|---|---|---|
| Huntington disease | 0.00010 | DCTN5, DNAH2, NDUFB10, PSMD13, HTT, DNAH9, COX5A, PSMB7, GRM5, TUBA1A, RB1CC1, UQCRFS1, NDUFV3, MAP3K5, COX8A, DNAH17, BDNF, ATG13, SDHB, SOD1, PSMA4, MAP3K10, ULK1, ACTR10, RCOR1 |
| Pathways of neurodegeneration | 0.00042 | CHRM3, DCTN5, DNAH2, NDUFB10, UBA7, PSMD13, HTT, DNAH9, COX5A, RYR3, PSMB7, ATXN3, GRM5, TUBA1A, RB1CC1, UQCRFS1, NDUFV3, MAP3K5, COX8A, WNT10A, DNAH17, BDNF, FZD6, ATG13, SDHB, SOD1, PSMA4, ALS2, BCL2, MAP3K10, ULK1, ACTR10 |
| RNA degradation | 0.00046 | CNOT4, BTG3, BTG2, BTG1, PNPT1, CNOT2, CNOT3, TOB1, LSM3, EDC3 |
| Amyotrophic lateral sclerosis | 0.00060 | DCTN5, DNAH2, NDUFB10, PSMD13, DNAH9, COX5A, PSMB7, TUBA1A, RB1CC1, TPR, UQCRFS1, NDUFV3, MAP3K5, COX8A, NUP210, DNAH17, NUP133, ATG13, SDHB, SOD1, PSMA4, ALS2, BCL2, PFN4, ULK1, ACTR10 |
| Longevity regulating pathway | 0.00333 | ATF2, PRKAA1, EHMT2, INSR, RB1CC1, ULK1, ATG13, ADIPOR2, IGF1R, SOD1 |
| Ribosome biogenesis in eukaryotes | 0.00502 | UTP15, TBL3, POP7, XPO1, TCOF1, POP4, AK6, GNL2, RPP38, MDN1 |
| Non-alcoholic fatty liver disease | 0.02237 | COX8A, PRKAA1, NDUFB10, INSR, UQCRFS1, NDUFV3, FOS, SDHB, ADIPOR2, COX5A, MAP3K5 |
| Alzheimer disease | 0.02310 | COX8A, CHRM3, WNT10A, NDUFB10, PSMD13, INSR, FZD6, LPL, ATG13, COX5A, SDHB, RYR3, PSMB7, GRM5, TUBA1A, PSMA4, RB1CC1, UQCRFS1, ULK1, NDUFV3, MAP3K5 |
| RNA transport | 0.03362 | PNN, DDX19A, POP7, XPO1, NUP210, NUP133, POP4, TPR, XPO5, RPP38, SNUPN, THOC6 |
| Mannose type O-glycan biosynthesis | 0.04637 | CRPPA, B3GAT1, POMT1 |

**Supplemental Figure 1: Differentially methylated genes are associated with neurodegenerative disease outcomes and the mTOR signaling pathway.** KEGG 2021 analysis was completed on all p-unadjusted DMR associated genes using Enrichr. We see differentially methylated genes are associated with neurodegenerative disease and the mTOR signaling pathway shows up as a top hypermethylated signaling pathway.

### Gene Ontology: Molecular Functions

| Top Upregulated GO Terms | P-Value | Top Downregulated GO Terms | P-Value |
|---|---|---|---|
| LIPOPOLYSACCHARIDE BINDING | 0.001 | CALCIUM CHANNEL ACTIVITY | 0.001 |
| METHIONINE SYNTHASE REDUCTASE ACTIVITY | 0.002 | POTASSIUM CHLORIDE SYMPORTER ACTIVITY | 0.001 |
| OXIDOREDUCTASE ACTIVITY OXIDIZING METAL IONS NAD OR NADP AS ACCEPTOR | 0.002 | G-PROTEIN COUPLED RECEPTOR KINASE ACTIVITY | 0.002 |
| PHOSPHATIDYLCHOLINE-STEROL O-ACYLTRANSFERASE ACTIVITY | 0.002 | NEUROTRANSMITTER RECEPTOR ACTIVITY | 0.002 |
| ZINC ION TRANSMEMBRANE TRANSPORTER ACTIVITY | 0.003 | RHODOPSIN KINASE ACTIVITY | 0.002 |
| ALPHA-1->6-FUCOSYLTRANSFERASE ACTIVITY | 0.004 | CATION CHLORIDE SYMPORTER ACTIVITY | 0.003 |
| GLYCOPROTEIN 6-ALPHA-L-FUCOSYLTRANSFERASE ACTIVITY | 0.004 | NEUROTRANSMITTER BINDING | 0.003 |
| CHROMATIN INSULATOR SEQUENCE BINDING | 0.006 | RAP GUANYL-NUCLEOTIDE EXCHANGE FACTOR ACTIVITY | 0.006 |
| IGE BINDING | 0.006 | TRANSLATION RELEASE FACTOR ACTIVITY | 0.006 |
| RNA POLYMERASE II CORE PROMOTER PROXIMAL REGISEQUENCE-SPECIFIC DNA BINDING ACTIVITY | 0.006 | TRANSLATION RELEASE FACTOR ACTIVITY CODON SPECIFIC | 0.006 |

### Gene Ontology: Cellular Components

| Top Upregulated GO Terms | P-Value | Top Downregulated GO Terms | P-Value |
|---|---|---|---|
| CILIUM AXONEME | 0.001 | SYNAPSE | 0.001 |
| CYTOSKELETAL PART | 0.004 | CELL JUNCTION | 0.002 |
| INHIBIN A COMPLEX | 0.009 | PROTEASOME REGULATORY PARTICLE BASE SUBCOMPLEX | 0.006 |
| LOW-DENSITY LIPOPROTEIN PARTICLE | 0.014 | GERMINAL VESICLE | 0.006 |
| ACTIVIN A COMPLEX | 0.017 | MICROTUBULE | 0.007 |
| F-ACTIN CAPPING PROTEIN COMPLEX | 0.017 | AXON | 0.008 |
| EXTRINSIC TO INTERNAL SIDE OF PLASMA MEMBRANE | 0.019 | CELL CORTEX | 0.009 |
| NUCLEAR PORE | 0.026 | NUCLEAR EUCHROMATIN | 0.010 |
| SMALL NUCLEAR RIBONUCLEOPROTEIN COMPLEX | 0.029 | MELANOSOME | 0.010 |
| CHYLOMICRON | 0.030 | NEURONAL CELL BODY | 0.010 |

**Supplemental Figure 2: Gene Ontology Molecular Functions and Cellular Components highlight further hypofunctioning neuronal systems and the important cell structures associated with Mn altered DEGs.** Further FGSEA analysis was conducted to assess Gene Ontology groupings of molecular functions and cellular components to gain additional insight into Mn-induced alterations of gene expression. The top down-and upregulated molecular functions reflect developmental Mn altered neuronal function (neurotransmitter receptor activity and binding). The top down-and upregulated cellular components reflect the genes altered by developmental Mn exposure primarily being related to neuronal cellular components (synapse, axon, cell cortex, neuronal cell body, axoneme).

### Top 10 Significant KEGG 2021 Terms

#### Downregulated

| Term | P-value | Overlap Genes |
|---|---|---|
| GABAergic synapse | 0.0001 | GABBR2, GABARAPL1, SLC12A5, GNG5, CACNA1A, PRKACB, GABRG2 |
| Morphine addiction | 0.0008 | GABBR2, GNG5, GRK4, CACNA1A, PRKACB, GABRG2 |
| Synaptic vesicle cycle | 0.0027 | SNAP25, STXBP1, ATP6V1B2, CACNA1A, STX3 |
| Ribosome | 0.0033 | RPS18, RPL34, RPL22, RPS20, RPL26, FAU, RPL17 |
| SNARE interactions in vesicular transport | 0.0075 | VAMP7, STX3, BET1 |
| Ubiquitin mediated proteolysis | 0.0076 | UBE2H, NEDD4, UBE2D3, UBE2O, TRIM37, UBE2K |
| Coronavirus disease | 0.0079 | MAPK10, RPS18, RPL34, RPL22, RPS20, RPL26, FAU, RPL17 |
| Cell adhesion molecules | 0.0099 | CADM3, CADM1, ITGA4, NRXN3, NRCAM, NCAM1 |
| Spliceosome | 0.0105 | HNRNPK, LSM6, ZMAT2, SNRNP70, PRPF3, BCAS2 |
| Nicotine addiction | 0.0128 | GRIN3B, CACNA1A, GABRG2 |

#### Upregulated

| Term | P-value | Overlap Genes |
|---|---|---|
| Thermogenesis | 0.00001 | NDUFA8, UQCRB, NDUFB6, NDUFA10, RPS6, ATP5MC2, TSC2, SMARCA4, DPF1, NDUFAB1, KRAS, UQCRC2, ATP5ME |
| Parkinson disease | 0.00002 | NDUFA8, UQCRB, NDUFB6, NDUFA10, ATP5MC2, PSMA7, PSMD6, TUBA1B, PSMB4, NDUFAB1, CALM3, UQCRC2, CALM1 |
| Alzheimer disease | 0.00002 | NDUFA8, UQCRB, NDUFB6, CSNK1A1, NDUFA10, ATP5MC2, ADAM10, PSMA7, PSMD6, TUBA1B, PSMB4, NDUFAB1, CALM3, KRAS, UQCRC2, CALM1 |
| Huntington disease | 0.00004 | NDUFA8, CREBBP, UQCRB, NDUFB6, NDUFA10, ATP5MC2, PSMA7, PSMD6, TUBA1B, PSMB4, SIN3A, NDUFAB1, UQCRC2, DNAL1 |
| Oxidative phosphorylation | 0.00005 | NDUFA8, UQCRB, NDUFB6, NDUFA10, NDUFAB1, ATP5MC2, UQCRC2, ATP6V1C1, ATP5ME |
| Pathways of neurodegeneration | 0.00014 | GABARAPL2, NDUFA8, UQCRB, NDUFB6, CSNK1A1, NDUFA10, ATP5MC2, PSMA7, PSMD6, TUBA1B, PSMB4, NDUFAB1, CALM3, KRAS, UQCRC2, CALM1, DNAL1 |
| Amyotrophic lateral sclerosis | 0.00027 | GABARAPL2, NDUFA8, UQCRB, NDUFB6, NDUFA10, ATP5MC2, PSMA7, PSMD6, TUBA1B, PSMB4, NDUFAB1, NUP54, UQCRC2, DNAL1 |
| Prion disease | 0.00082 | NDUFA8, PSMD6, TUBA1B, PSMB4, UQCRB, NDUFB6, NDUFAB1, NDUFA10, ATP5MC2, UQCRC2, PSMA7 |
| Phosphatidylinositol signaling system | 0.00155 | DGKG, MTMR2, INPP1, PI4KA, CALM3, CALM1 |
| Non-alcoholic fatty liver disease | 0.00388 | NDUFA8, UQCRB, NDUFB6, NDUFA10, NDUFAB1, LEPR, UQCRC2 |

**Supplemental Figure 3: KEGG 2021 Analysis confirms that the genes altered by developmental Mn are associated with hypofunctioning neuronal systems and neurodegenerative disease outcomes.** KEGG 2021 analysis was conducted on p-unadjusted significant DEGs using Enrichr. Downregulated KEGG terms show neuronal function deficits (GABAergic synapse and synaptic vesicle cycle) and upregulated genes showing significant relationship to neurological disorders and neurodegenerative diseases that are associated with proinflammatory mechanisms (Parkinson's, Huntington's, and Alzheimer's disease). The overlap genes that most contributed to each gene terms significance are listed with several of them being from the 27 p-adjusted genes of Figure 4A, such as Prkacb and Crebbp.

| GO Biological Processes Terms | Expression Direction | Functional Category |
|---|---|---|
| Regulation of immune response | Upregulated | Inflammation |
| Apoptotic signaling pathway | Upregulated | Inflammation |
| Astrocyte fate commitment | Upregulated | Inflammation |
| Regulation of gene expression, epigenetic | Upregulated | Epigenetics |
| Regulation of DNA methylation | Upregulated | Epigenetics |
| Regulation of histone methylation | Upregulated | Epigenetics |
| Negative regulation of Wnt receptor signaling pathway | Upregulated | Cell Development |
| TOR signaling | Upregulated | Cell Development |
| Positive regulation of catecholamine secretion | Downregulated | Neuronal Function |
| Neuronal action potential propagation | Downregulated | Neuronal Function |
| Synaptic transmission | Downregulated | Neuronal Function |
| Learning | Downregulated | Neuronal Function |

**Supplemental Figure 4: Notable GO upregulated and downregulated terms to our Mn neurotoxicity epigenetic and catecholaminergic phenotype.** $p < 0.05$ significant GO biological processes terms, their differential gene expression fold change direction, and their corresponding functional category (ie. inflammation, epigenetic, cell development, and neuronal function from the differential expression analysis.

### Significant Gene Ontology Pathways

**A** Downregulated

#### Biological Process

| Gene Ontology Pathway | P-Value |
|---|---|
| Calcium ion transmembrane transport | 0.029 |
| Protein insertion into membrane | 0.029 |
| Regulation of calcium ion transport | 0.029 |
| Synaptic transmission | 0.029 |
| Desensitization of g-protein coupled receptor protein signaling pathway | 0.039 |
| G-protein coupled receptor internalization | 0.039 |
| Ion transport | 0.039 |
| Receptor internalization | 0.039 |

#### Molecular Functions

| Gene Ontology Pathway | P-Value |
|---|---|
| Calcium channel activity | 0.029 |
| Cation channel activity | 0.029 |
| Extracellular-glutamate-gated ion channel activity | 0.029 |
| Glycine binding | 0.029 |
| Ionotropic glutamate receptor activity | 0.029 |
| Neurotransmitter binding | 0.029 |
| Neurotransmitter receptor activity | 0.029 |
| G-protein coupled receptor kinase activity | 0.039 |
| Rhodopsin kinase activity | 0.039 |

#### Cellular Components

| Gene Ontology Pathway | P-Value |
|---|---|
| Neuronal cell body | 0.009 |
| Postsynaptic membrane | 0.019 |
| Cell cortex | 0.022 |
| N-methyl-d-aspartate selective glutamate receptor complex | 0.029 |
| Outer membrane-bounded periplasmic space | 0.029 |

**B** Upregulated

#### Biological Process

| Gene Ontology Pathway | P-Value |
|---|---|
| Positive regulation of gene expression | 0.029484 |
| Protein phosphorylation | 0.034398 |
| Transmembrane transport | 0.039669 |
| Actin cytoskeleton organization | 0.049524 |
| Cytokine-mediated signaling pathway | 0.049524 |
| Negative regulation of neuron apoptotic process | 0.049524 |
| Positive regulation of MAP kinase activity | 0.049524 |
| Positive regulation of NF-KAPPAb transcription factor activity | 0.049524 |
| Positive regulation of nitric-oxide synthase activity | 0.049524 |
| Positive regulation of protein phosphorylation | 0.049524 |
| Positive regulation of RAC protein signal transduction | 0.049524 |
| Regulation of long-term neuronal synaptic plasticity | 0.049524 |
| Regulation of synaptic transmission gabaergic | 0.049524 |
| Social behavior | 0.049524 |
| Visual learning | 0.049524 |

#### Molecular Functions

| Gene Ontology Pathway | P-Value |
|---|---|
| GTPase activator activity | 0.001653 |
| Actin binding | 0.028571 |
| Protein kinase activity | 0.034398 |
| Protein serine threonine kinase activity | 0.034398 |
| Transferase activity transferring phosphorus-containing groups | 0.034398 |
| GMP binding | 0.049524 |
| LRR domain binding | 0.049524 |

#### Cellular Components

| Gene Ontology Pathway | P-Value |
|---|---|
| Cell projection | 0.028571 |
| Membrane raft | 0.039669 |

**Supplemental Figure 5: GO analysis of the 155 Genes both differentially methylated and differentially expressed highlight hypofunctioning neuronal biological, molecular, and cellular components.** GO biological function (BF), molecular function (MF), and cellular components (CC) analysis of the 155 Genes both differentially methylated and differentially expressed highlight similar inflammatory (cytokine-mediated signaling pathway, regulation of neuron apoptotic process, positive regulation of nitric-oxide synthase activity), epigenetic (positive regulation of gene expression), cell developmental (positive regulation of RAC protein signal transduction and actin cytoskeleton organization), and hypofunctioning neuronal pathways (synaptic transmission, neurotransmitter binding, G-protein coupled receptor internalization, regulation of long-term neuronal synaptic plasticity) to those that have been previously discussed. Several of these BP and MF can be linked to mTOR function, such as the protein phosphorylation and protein kinase activity. It is also interesting for CC that neuronal cell body, postsynaptic membrane, and cell cortex were the three main downregulated pathways highlighting that Mn neurotoxicity in these 155 both DMR and DEG genes is highly related to neuronal function.

### Top 10 Significant KEGG 2021 Terms

#### Downregulated

| Term | P-value | Overlap Genes |
|---|---|---|
| Spinocerebellar ataxia | 0.002 | MAPK10, GRIN3B, OPA1, CACNA1A |
| GABAergic synapse | 0.005 | GABBR2, GABARAPL1, CACNA1A |
| Morphine addiction | 0.005 | GABBR2, GRK4, CACNA1A |
| Nicotine addiction | 0.010 | GRIN3B, CACNA1A |
| Type II diabetes mellitus | 0.013 | MAPK10, CACNA1A |
| Fc epsilon RI signaling pathway | 0.028 | MAPK10, FYN |
| Mitophagy | 0.028 | MAPK10, GABARAPL1 |
| NOD-like receptor signaling pathway | 0.033 | MAPK10, GABARAPL1, PKN1 |
| RNA transport | 0.036 | XPO1, EEF1A2, NUP58 |
| RNA degradation | 0.038 | CNOT4, LSM6 |

#### Upregulated

| Term | P-value | Overlap Genes |
|---|---|---|
| mTOR signaling pathway | 0.003 | ACTR2, AFDN, TUBA1B, ARPC1A |
| Tight junction | 0.004 | TUBA1B, GNA11, KRAS |
| Gap junction | 0.004 | PDPK1, TSC2, KRAS |
| Choline metabolism in cancer | 0.006 | XRCC1, LIG3 |
| Base excision repair | 0.007 | PDPK1, KRAS |
| Aldosterone-regulated sodium reabsorption | 0.009 | RAB2A, PDPK1, TSC2 |
| AMPK signaling pathway | 0.011 | PDPK1, TSC2, KRAS |
| Thyroid hormone signaling pathway | 0.011 | PDPK1, TSC2, KRAS |
| Insulin signaling pathway | 0.015 | PDPK1, TSC2, KRAS |
| Autophagy | 0.015 | ACTR2, AFDN, TUBA1B, ARPC1A |

**Supplemental Figure 6: KEGG analysis of the 155 Genes both differentially methylated and differentially expressed highlight mTOR signaling pathway as the most significant upregulated KEGG term and additional hypofunctioning neuronal function.** KEGG 2021 was performed in Enrichr on the 155 Genes (78 Upregulated/77 Downregulated) that were DMR and DEG. To further investigate the mechanistic insights of the 155 Genes both differentially methylated and differentially expressed, KEGG analysis was done and found the mTOR signaling pathway as the most significant upregulated KEGG term ($p < 0.05$). Additionally, KEGG analysis showed several other hypofunctioning neuronal functions, such as downregulated Spinocerebellar ataxia, GABAergic synapse, and RNA degradation and upregulated tight junction, gap junction, base excision repair, and autophagy ($p < 0.05$).

| Gene | Function | Biological Function Category |
|---|---|---|
| DnaJ Heat Shock Protein Family (Hsp40) Member C5 (DNAJC5) | Regulates ATPase activity. Acts as a co-chaperone for the SNARE protein SNAP-25 and involved in calcium-dependent neurotransmitter release. | Neuronal and Cellular Function |
| Quaking (QKI) | Regulates pre-mRNA splicing, export of mRNAs from the nucleus, protein translation, and mRNA stability. Is involved in myelinization and oligodendrocyte differentiation connecting mTOR and Wnt signaling pathways. | Cellular Differentiation and Proliferation |
| Leucine Rich Repeat Containing 4 (LRRC4) | Synaptic adhesion protein. Regulates the formation of excitatory synapses through the recruitment of pre-and-postsynaptic proteins. Organize the lamina/pathway-specific differentiation of dendrites. | Neuronal and Cellular Function |
| Inner Membrane Mitochondrial Protein (IMMT) | Enables RNA binding activity. Involved in cristae formation. Located in mitochondrial inner membrane. Part of MICOS complex. | Oxidative Stress and Inflammation |
| Coiled-Coil-Helix-Coiled-Coil-Helix Domain Containing 6 (CHCHD6) | Involved in cellular response to DNA damage stimulus and cristae formation. Located in cytosol and mitochondrial inner membrane. Part of the MICOS complex. | Epigenetics and DNA Damage |
| Gamma-Aminobutyric Acid Type B Receptor Subunit 2 (GABBR2) | GABA-B receptors inhibit neuronal activity through G protein-coupled second-messenger systems, which regulate the release of neurotransmitters, and the activity of ion channels and adenylyl cyclase. | Neuronal and Cellular Function |
| Protein Kinase CAMP-Activated Catalytic Subunit Beta (PRKACB) | Mediates cAMP-dependent signaling triggered by receptor binding to GPCRs that is important for activating QKI and DARPP-32 signaling. | Neuronal and Cellular Function |
| Ubiquitin C-Terminal Hydrolase L3 (UCHL3) | Deubiquitinating enzyme that controls levels of cellular ubiquitin, May be involved in tauopathy and synucleinopathy. | Neuronal and Cellular Function |

**Supplemental Figure 7: DMR and DEG integration with IPA elucidates important gene regulators of our Mn neurotoxicity phenotype.** Key genes determined to be upstream and causal regulators by Ingenuity Pathway Analysis (IPA). A brief definition of their function and biological relevance to our developmental hypothesis are displayed.

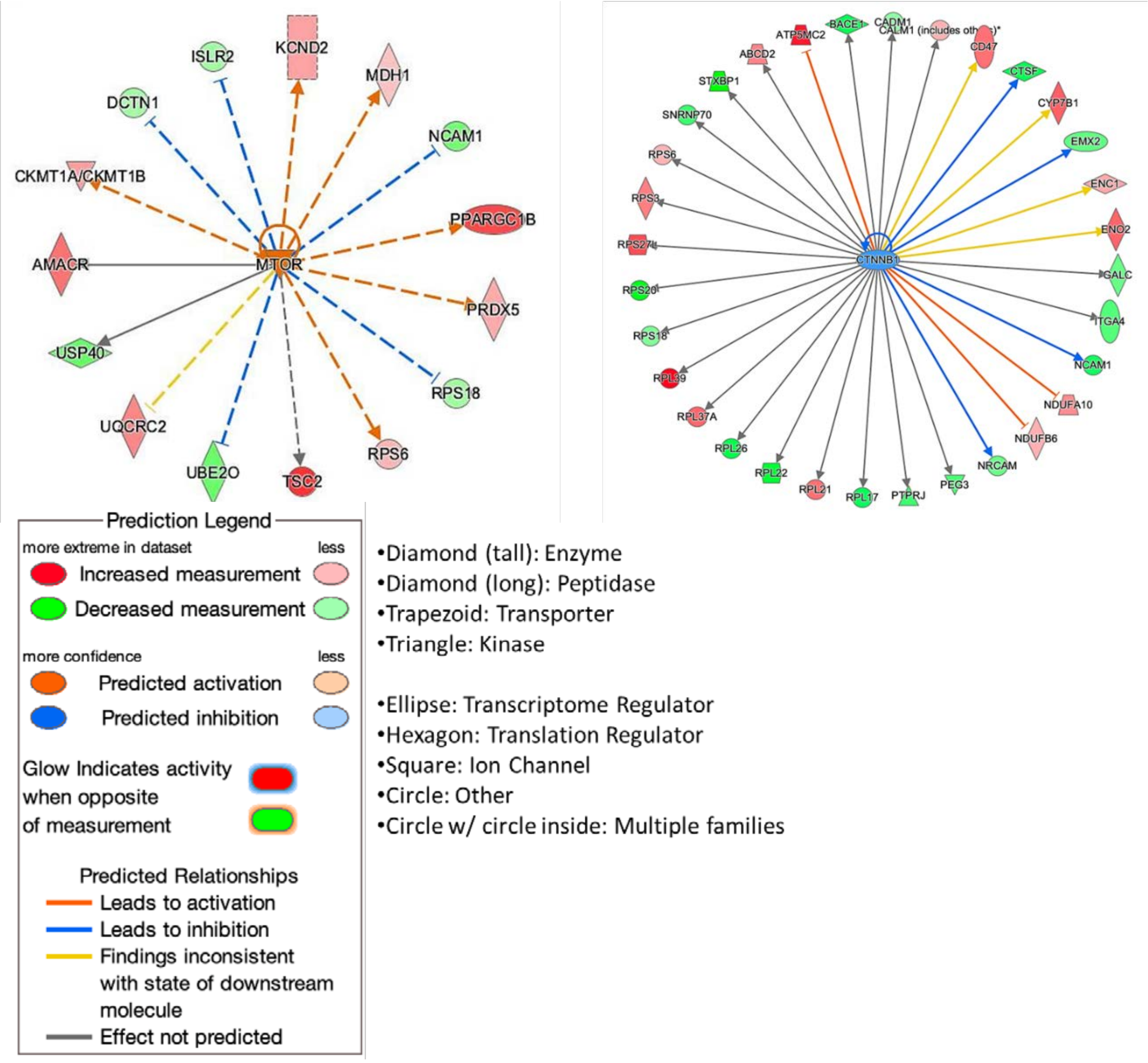


**Supplemental Figure 8: DMR and DEG integration further demonstrates the mechanistic importance of Wnt and mTOR signaling pathway.** Regulator analysis of mTOR and Wnt within the current data set shows both of their significant DEG networks determined by IPA. The number of genes they serve as upstream regulators for further emphasizes the importance of these two pathways as being master contributors to our proposed Mn neurotoxicity molecular and behavioral phenotype.

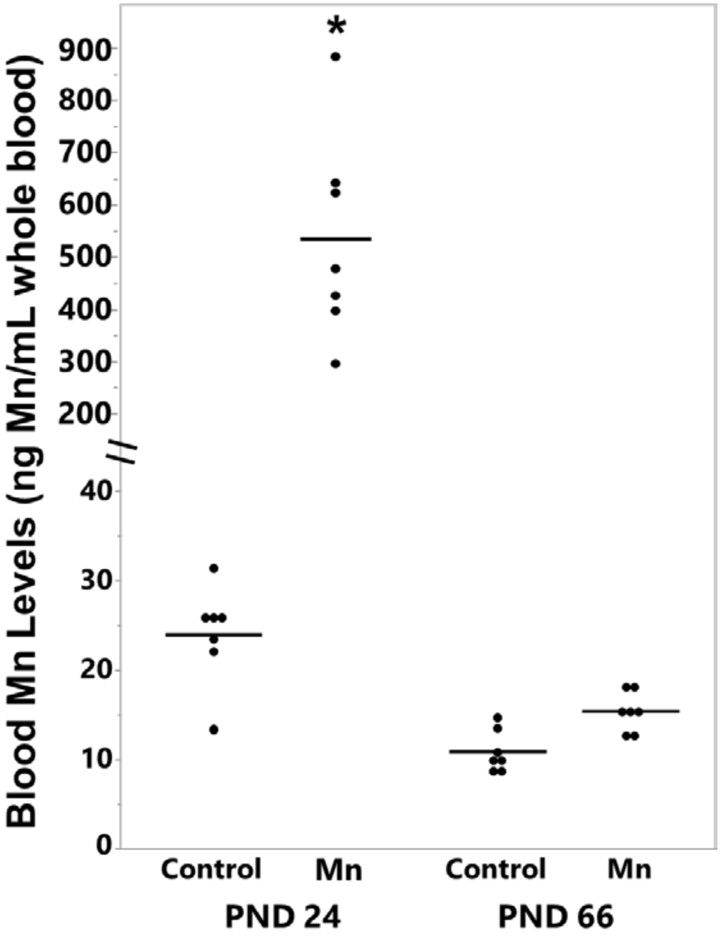


**Supplemental Figure 9: Developmental Mn exposure increased blood manganese levels and these levels normalize by PND 66.** To confirm that our developmental exposure regimen increased Mn levels in our experimental rodents to environmentally relevant levels and thus, induces our resulting behavioral and molecular phenotypes, we measured Mn levels in the blood of PND 24 animals via ICP-MS. Our developmental Mn exposure regimen significantly increased blood Mn levels of PND 24 Mn exposed rats compared to PND 24 controls ($p < 0.001$). To determine that the resulting Mn-induced behavioral and molecular phenotypes are due to lasting effects of neurochemical changes of developmental Mn exposure and not due to active elevated Mn exposure we assessed whether blood Mn levels are reduced 45 days following the last Mn dose at PND 66. The blood Mn levels of animals exposed to Mn during PND 1-21 are reduced to control levels 45 days after Mn exposure has ceased ($p < 0.001$). The data shows that the following behavioral and molecular phenotypes associated with Mn exposure are due to lasting neurochemical changes caused by developmental Mn exposure and not due to active elevated Mn exposure at time of behavioral testing and molecular measurements are assessed.
